# The alternative sigma factor RpoD4 pulses at division and links the clock and the cell cycle in *Synechococcus elongatus*

**DOI:** 10.1101/2023.06.28.546855

**Authors:** Chao Ye, Teresa Saez, Arijit K. Das, Bruno M.C. Martins, James C.W. Locke

**Affiliations:** Sainsbury Laboratory, University of Cambridge, Cambridge, UK; School of Life Sciences, University of Warwick, Coventry, UK

## Abstract

The cyanobacterial circadian clock is an extensive regulator of gene expression, generating a 24-hour rhythm in the majority of genes in *Synechococcus elongatus* PCC *7942*. This raises the question of how the cyanobacterial clock regulates, and receives input from, the diverse cellular processes it controls. Alternative sigma factors, which bind to the core RNA polymerase enzyme and direct the synthesis of specific sets of transcripts, are one mechanism for the clock to set the time of its outputs. In this work, we use single-cell time-lapse microscopy to reveal the transcriptional dynamics of RpoD4, an alternative sigma factor that was previously reported as either arrhythmic or circadian. We find instead that RpoD4 pulses at cell division, dynamics missed by previous bulk averaging. The circadian clock modulates the amplitude of RpoD4 expression pulses, as well as the timing of pulses through its control of division timing. In turn, a *rpoD4* mutation causes a reduction in clock period. Further, a *rpoD4* mutant results in smaller cell size, and increasing expression of RpoD4 results in larger cell sizes in a dose-response manner, allowing tunable control of cell size. Thus, our single-cell analysis has revealed pulsing gene expression dynamics in cyanobacteria, linking the clock, RpoD4 and the cell cycle.

## Introduction

Circadian clocks are biological pacemakers that oscillate with a circa 24 h period (Doherty & Kay, 2010; Golden & Canales, 2003; Hurley *et al*, 2016; Paranjpe & Sharma, 2005). Many organisms across different taxa have evolved circadian clocks, as clocks allow anticipation of daily environmental changes caused by the rotation of the earth. Although the molecular mechanisms underlying circadian clocks have been studied extensively, how circadian clocks couple to other biological processes in individual cells and what the physiological consequences of such coupling are remains a topic of intense research interest (Bieler *et al*, 2014; Feillet *et al*, 2015; Gould *et al*, 2018; Lambert *et al*, 2016; Liao & Rust, 2021; Martins *et al*, 2018).

*Synechococcus elongatus PCC 7942* (*S. elongatus*) is an ideal model for studying the coupling of the clock to cellular processes in individual cells. The clock is well characterised (Cohen & Golden, 2015; Johnson & Egli, 2014), and can be monitored in individual cells over time (Chabot *et al*, 2007). The core network consists of just three proteins (KaiA, KaiB, and KaiC) that generate a 24-h oscillation in KaiC phosphorylation. This core oscillator controls the activity of the master regulator RpaA, which goes on to orchestrate global gene expression (Markson *et al*, 2013). In *S. elongatus*, most pathways are modulated by the circadian clock (Ito *et al*, 2009; Vijayan *et al*, 2009). These include photosynthesis (Martins *et al*, 2016; Welkie *et al*, 2018), cell division, and growth (Dong *et al*, 2010; Mori *et al*, 1996; Martins *et al*, 2018; Yang *et al*, 2010), but also stress response (Hanaoka & Tanaka, 2008; Hanaoka *et al*, 2012; Kobayashi *et al*, 2017; Moronta-Barrios *et al*, 2012; Seki *et al*, 2007), metabolism (Johnson & Egli, 2014; Pattanayak & Rust, 2014) and phototaxis (Yang *et al*, 2018). One proposed mechanism to allow the clock to regulate such a range of processes is through a set of group 2 alternative sigma factors (Markson *et al*, 2013; Fleming & O’Shea, 2018).

The first systematic study of gene expression dynamics of group 2 sigma factors was conducted by Nair *et al*. (Nair *et al*, 2002), who used luciferase reporters to demonstrate that *rpoD2*, *rpoD3* and *sigC* are transcribed with circadian rhythms at the population level. Although the authors reported that the RpoD4 protein level oscillates with a circadian rhythm and peaks at a circadian time (CT) of 4 h, the transcriptional dynamics of *rpoD4* were not reported. Fleming and O’Shea (Fleming & O’Shea 2018) went further and found that four RpaA-dependent sigma factors—RpoD2, RpoD6, SigC, and SigF2—are sequentially activated downstream of RpaA and are required for proper expression of circadian mRNAs. SigC has also been proposed to form an oscillatory feedforward loop with the clock and downstream targets, which can allow frequency doubling of the circadian oscillator (Martins *et al*, 2016).

RpoD4 is one of two group 2 sigma factors not regulated by RpaA. Previous microarray studies have characterised it either as circadian (Vijayan *et al*, 2009) or arrhythmic (Ito *et al*, 2009). In this work, we set out to understand the activation dynamics of RpoD4 in individual cells using single-cell timelapse microscopy. We found that RpoD4 expression is pulsatile, with peaks occurring at cell division. The circadian clock modulates the amplitude of the pulses, causing the largest pulses to occur around subjective dusk. RpoD4 expression also feeds back to the clock, as a *rpoD4* deletion causes a reduction in clock period and over-expression of RpoD4 increases clock period. Finally, *rpoD4* deletion causes a reduction in cell size in both WT and clock deletion strains, with overexpression of RpoD4 increasing cell size. Thus, our work has revealed pulsing dynamics in cyanobacteria, with subtle interconnections between the clock, RpoD4, and the cell cycle.

## Results

### *rpoD4* expression pulses at cell division

To track expression of *rpoD4* in individual cells, we first constructed a transcriptional reporter strain containing a chromosomally integrated fluorescent reporter for *rpoD4*, P*_rpoD4_*-EYFP-LVA. We then used time-lapse microscopy to examine P*_rpoD4_*-EYFP expression in individual cells grown on agarose pads under constant light conditions (c.a. 18 μE m^-2^ s^-1^) (**Materials and methods**). Before the start of the movie, cells were entrained by 12:12 h square wave light-dark (LD) cycles before being released into constant light (LL). From our time-lapse movies, we extracted and compared the single-cell time traces of cell length and P*_rpoD4_*-EYFP expression dynamics, and found that the expression of *rpoD4* pulses at cell division, rather than showing circadian oscillations (**Fig. 1A-C**).

**Figure 1.**
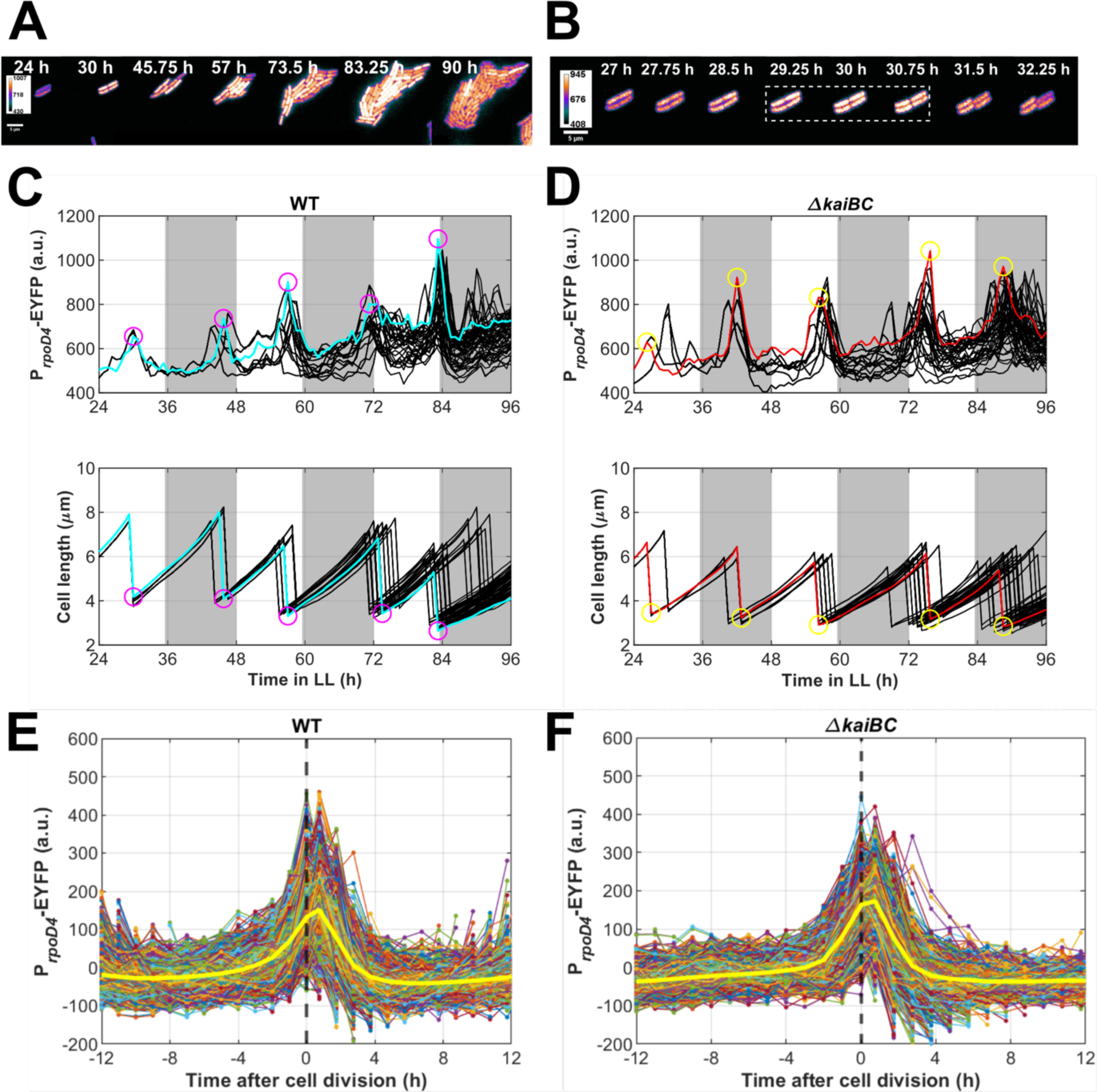
*rpoD4* expression peaks at cell division. (**A**) Montage of fluorescence microscopy images of WT P*_rpoD4_*-EYFP-LVA cells grown under constant light (c.a. 18 μE m^-2^ s^-1^). The heatmap indicates the YFP fluorescence intensity range. (**B**) Eight consecutive frames from the same time-lapse movie in (**A**) demonstrating the expression of P*_rpoD4_*-EYFP pulses during cell division. The dashed white box highlights cytokinesis. (**C**) Profiles of P*_rpoD4_*-EYFP expression and cell length in WT and (**D**) *ΔkaiBC* backgrounds grown under constant light (LL). Each black line shows a lineage, and the coloured lines are representative lineages. Note the peaks of P*_rpoD4_*-EYFP expression (prominence ≥ 100 a.u.) and the troughs in the cell length profile (corresponding to cell division events) are correlated as highlighted by coloured circles. The white background represents the subjective day, and the grey shades represent the subjective night. (**E**) Single-cell expression dynamics of detrended P*_rpoD4_*-EYFP in WT and (**F**) *ΔkaiBC* backgrounds plotted against time centred at cell division (t = 0 h). Lines represent individual cells with complete cell cycles. The solid yellow line represents the mean. The dashed black line indicates t = 0 h. For both WT and *ΔkaiBC*, expression of P*_rpoD4_*-EYFP peaks at or close to cell division. 19 WT movies and 18 *ΔkaiBC* movies from 2 independent experiments were collected.

To test whether *rpoD4* expression dynamics required the circadian clock, we repeated the experiment with the same P*_rpoD4_*-EYFP-LVA reporter, but in a clock-deletion (*ΔkaiBC*) strain. We found the expression of *rpoD4* still pulses and peaks at cell division without the clock (**Fig. 1D**). By pooling the time traces of P*_rpoD4_*-EYFP expression dynamics from all time-lapse movies and plotting them against the time around cell division (with time of division = 0 h), we found for both WT and *ΔkaiBC* reporter strains, the expression of *rpoD4* peaks at around t = 0.75 h (**Fig. 1E&F**).

As transcriptional dynamics may not be translated into protein dynamics, we next examined the post-translational dynamics of the RpoD4 protein, by constructing a translational reporter, P*_rpoD4_*-RpoD4-EYFP. Similar to the transcriptional reporter, the levels of the RpoD4-EYFP protein fusion also pulsed at cell division in both the WT and *ΔkaiBC* background (**Fig. S1A-D**), although with a lower signal-to-noise ratio compared to the transcriptional reporter. By aligning the time traces of P*_rpoD4_*-RpoD4-EYFP expression from all lineages relative to time of division, we found for both WT and *ΔkaiBC* reporter strains, the expression of RpoD4-EYFP peaks at around t = 0.75 h, (**Fig. S1E&F**), as for the P*_rpoD4_*-EYFP-LVA reporter. Taken together, our results suggest the expression of *rpoD4* pulses at cell division and the pulsing does not require a functional clock.

### The circadian clock modulates RpoD4 pulse amplitude

Given that the circadian clock was not required for RpoD4 pulsing, we tested whether there was any effect of the clock on RpoD4 expression or the characteristics of the observed pulses. We found that the averaged expression of P*_rpoD4_*-EYFP in WT displayed weak circadian oscillations that peak towards the end of the subjective day **(Fig. S2A)**. Circadian rhythmicity analysis on detrended mean P*_rpoD4_*-EYFP expression revealed that the WT strain, but not the *ΔkaiBC* strain, displayed circadian oscillations. The period analysis revealed that the detrended mean P*_rpoD4_*-EYFP expression in WT has a period of 24.7 h and an acrophase (the peak of a cycle) of CT 7 h (**Fig. S2B**), just a few hours before subjective dusk. This acrophase value falls within the time window where, due to clock regulation of cell division, most divisions are observed, which is also before subjective dusk (peak at CT 8.25 h, **Fig. S2E**) (Mori *et al*, 1996; Martins *et al*, 2018). Apparent circadian rhythms in *rpoD4* expression in bulk (Vijayan *et al*, 2009) are therefore, or at least in part, caused by rhythms in the timing of cell division events, with higher average expression expected to be observed in windows with more such events. The *ΔkaiBC* strain displayed an apparent shorter period rhythm of *rpoD4* expression of about 16 h (**Fig S2D**). This period is close to the mean cell cycle duration (15.9 h) and is therefore an artefact caused by a bias towards division events at certain time windows introduced by a ‘founder effect’ (colonies start from one cell) compounded by LD cycle division resetting (all cultures, both WT and *ΔkaiBC* were similarly entrained before the start of the experiment).

To examine the effect of the circadian clock on *rpoD4* expression peaks, we extracted unique peaks from detrended P*_rpoD4_*-EYFP expression profiles of all lineages. We found that peak height and peak width of P*_rpoD4_*-EYFP expression show prominent circadian rhythms in the WT strain **(Fig. 2A&C)**. On the other hand, deleting *kaiBC* abolishes the circadian rhythm of peak height and width of P*_rpoD4_*-EYFP expression **(Fig. 2B&D)**. These results suggest that the RpoD4 pulse height is modulated by the circadian clock and peaks around the subjective dusk.

**Figure 2.**
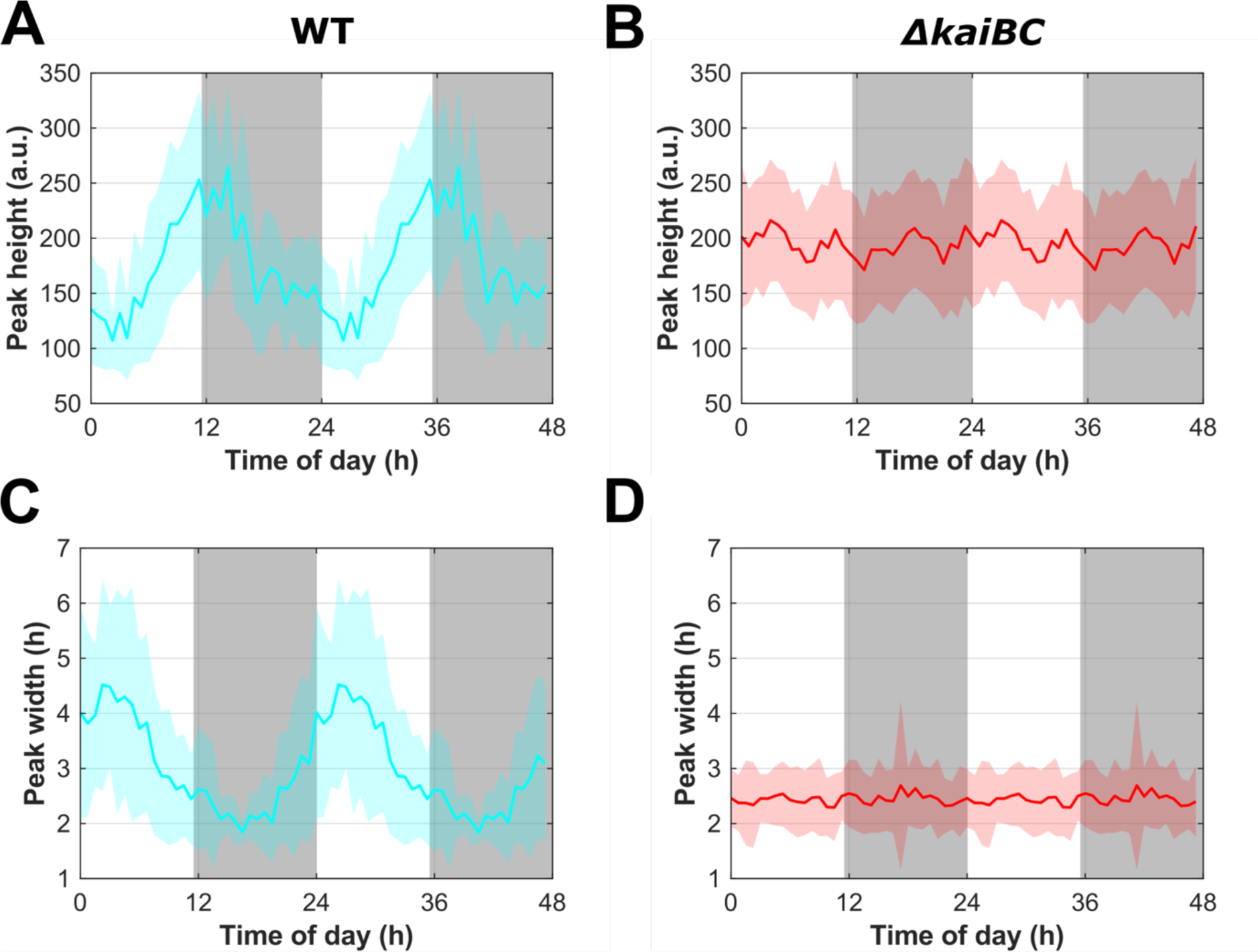
The circadian clock modulates the height and width of *rpoD4* expression peaks. (**A**) The averaged peak height and (**C**) width of *rpoD4* expression in the WT strain, depicted for two cycles to highlight periodicity, display circadian rhythms (solid cyan line, with cyan shades representing one standard deviation from the mean). 1870 peaks extracted from 2373 detrended lineage profiles from two independent experiments were analysed. In contrast, (**B**) the mean peak height and (**D**) width of *rpoD4* expression in the *ΔkaiBC* strain do not show any circadian rhythm (solid red line, with red shades representing one standard deviation from the mean). 2287 peaks extracted from 2775 detrended lineage profiles from 2 independent experiments were analysed.

We next investigated further the structure of *rpoD4* promoter to identify regions within the promoter required for pulsing. We constructed P*_rpoD4_*-EYFP-LVA reporters with varied promoter length ranging from 60 to 500 bp upstream of the *rpoD4* start codon, and a truncated promoter with 300-50 bp upstream of the *rpoD4* start codon. By analysing the single-cell time traces of P*_rpoD4_*-EYFP expression, we found that strains with promoters P*_rpoD4_*(500), P*_rpoD4_*(200) and P*_rpoD4_*(100) (the numbers in brackets represent the promoter length) all show pulsing at cell division **(Fig. S3B-D)**. However, when the promoter length is reduced to 60 bp, the expression of P*_rpoD4_*-EYFP becomes less robust, with many peaks, especially the ones that should occur near subjective dawn, not occurring (**Fig. S3E**). Furthermore, when using the truncated promoter P*_rpoD4_*(300-50), the expression of P*_rpoD4_*-EYFP can no longer be detected **(Fig. S3F)**, and the YFP fluorescence from this strain is similar to the WT background **(Fig. S3G)**. The results suggest that P*_rpoD4_*(100) is the minimal promoter for driving robust P*_rpoD4_*-EYFP-LVA expression, thus the cis-elements of the *rpoD4* promoter should reside within the 100 bp upstream of the *rpoD4* start codon.

### RpoD4 levels modulate clock period

We next investigated the physiological roles of RpoD4, given its pulsatile single-cell gene expression dynamics. We first examined whether RpoD4 affected the circadian clock, as previously Nair *et al*. (Nair *et al*, 2002) used luciferase reporters to show that overexpression of *rpoD4* results in an approximate 2 h period lengthening of the clock. In their study, deletion of *rpoD4* did not cause a significant change in clock period, although it did cause a shift in the phase of clock outputs. We tested whether with our time-lapse movies we could detect subtle effects of a *rpoD4* deletion on clock period potentially missed by bulk luciferase measurements. We first examined the expression dynamics of the *kaiBC* operon promoter, using a P*_kaiBC_*-EYFP-fsLVA reporter in WT and *ΔrpoD4* backgrounds under constant light conditions (c.a. 15 μE m^-2^ s^-1^). It should be noted a frame-shifted SsrA-LVA (fsLVA) degradation tag was identified in the widely used plasmid pJRC35 (Chabot *et al*, 2007; Gan & O’Shea, 2017; Pattanayak *et al*, 2015; Teng *et al*, 2013; Yang *et al*, 2010), which encodes the *S. elongatus* circadian clock reporter P*_kaiBC_*-EYFP-LVA (Chabot *et al*, 2007). By sequencing pJRC35, we found the LVA tag in pJRC35 contains a frameshift mutation, leading to a truncated degradation tag, causing the reporter to be more stable with a measured degradation half-life of 5.6 h (Chabot *et al*, 2007). We applied colony-level analysis (**Materials and methods**) to generate time traces of mean P*_kaiBC_*-EYFP expression from individual movies (**Fig. 3A**). After detrending the data, we performed period analysis of the data using the FFT-NLLS method (with period window constrained to 22-28 h) from the BioDare2 suite (Zielinski *et al*, 2014). As shown in **Fig. 3B**, deletion of *rpoD4* caused a slight 0.4 h reduction in the period of *kaiBC* expression (p = 2.97e-5, Wilcoxon rank-sum test) under constant light conditions (c.a. 15 μE m^-2^ s^-1^). To test whether this observation was specific to our reporter and growth conditions, we examined a clock reporter with a different degradation tag, P*_kaiBC_*-EYFP-ASV, under higher light intensity (c.a. 35 μE m^-2^ s^-1^). We performed coarse colony-level analysis followed by period analysis of the detrended data (**Fig. S4A**) again using FFT-NLLS in BioDare2. Deletion of *rpoD4* again leads to a small 0.3 h reduction in the period of *kaiBC* expression (p = 2.8e-5, Wilcoxon rank-sum test) (**Fig. S4B**). Taken together, these results indicate deletion of *rpoD4* causes a subtle but statistically significant reduction of the period of *kaiBC* expression across different light levels.

**Figure 3.**
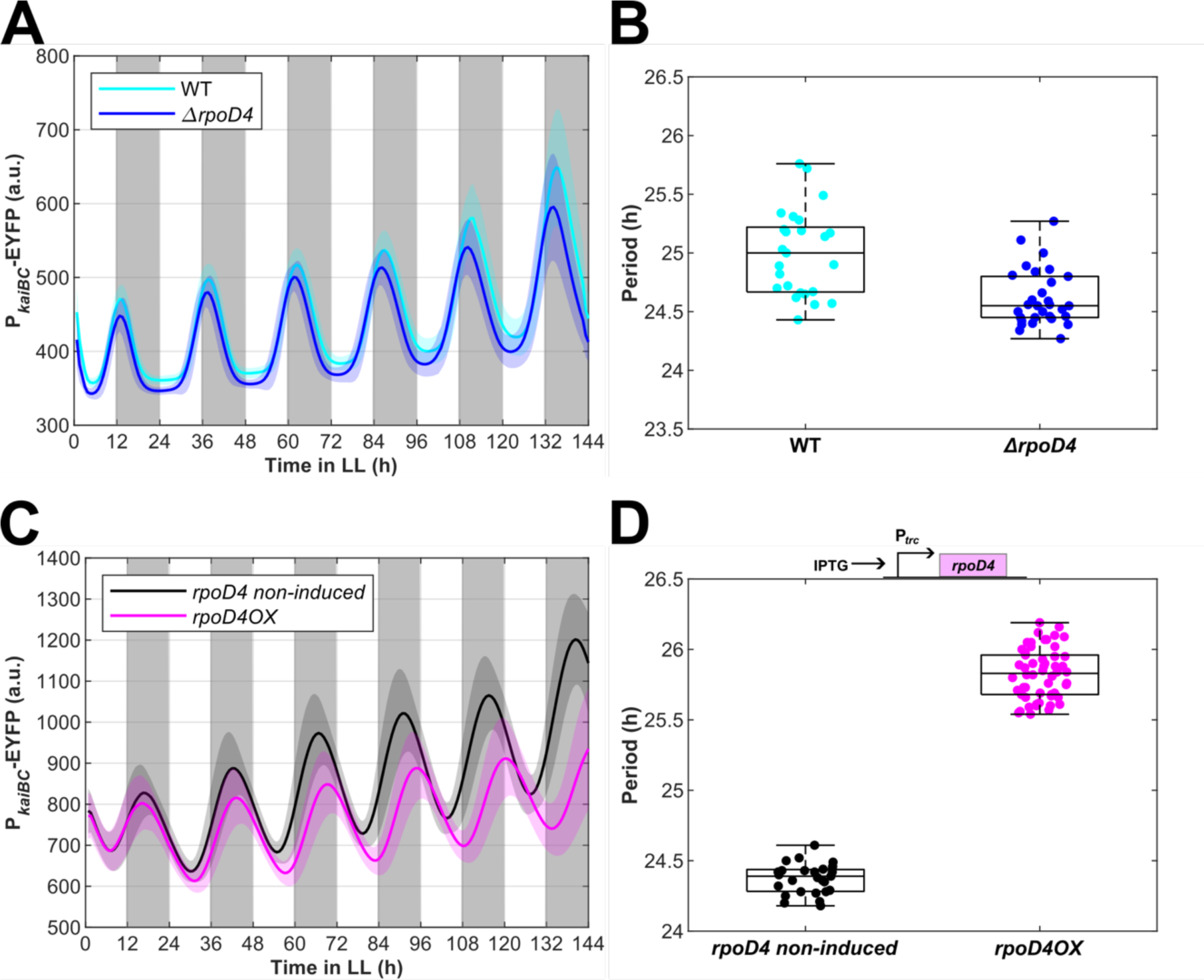
Deletion and overexpression of *rpoD4* modulate the period of P*_kaiBC_*-EYFP expression. (**A**) Time trace of mean P*_kaiBC_*-EYFP-fsLVA expression of WT (cyan line, averaged from 25 movies) and *ΔrpoD4* (blue line, averaged from 30 movies) strains grown under constant light conditions (c.a. 15 μE m^-2^ s^-1^). Data were collected from two independent experiments. The cyan and blue shades represent one standard deviation from the mean (**B**) Box plots of the period of detrended mean P*_kaiBC_*-EYFP-fsLVA expression in WT (cyan dots, n = 25, median ≈ 25 h) and *ΔrpoD4* (blue dots, n = 30, median ≈ 24.55 h) movies. The difference is statistically significant (p = 2.97e-5). (**C**) Time trace of mean P*_kaiBC_*-EYFP-fsLVA expression in cells carrying non-induced P*_trc_*-*rpoD4* (black line, averaged from 27 movies) and 100 mM IPTG induced P*_trc_-rpoD4* (abbreviated as *rpoD4OX*, magenta line, averaged from 54 movies) strains (**D**) Box plots of the period of detrended mean P*_kaiBC_*-EYFP-fsLVA expression in non-induced P*_trc_*-*rpoD4* (black dots, n = 27, median ≈ 24.39 h) and *rpoD4OX* (magenta dots, n = 54, median ≈ 25.83 h) movies. The difference is statistically significant (p = 2.82e-13). Diagram on top of the (**D**) depicts the inducible construct. Wilcoxon rank-sum test is used for all statistical tests in this figure.

We then checked the effects of *rpoD4* deletion on the dusk expressed gene *sigC*, and observed a similarly small but statistically significant period reduction in *sigC* expression (WT median ≈ 24.34 h, *ΔrpoD4* median ≈ 24.14 h, p = 4.23e-6, Wilcoxon rank-sum test) (**Fig. S4C**). Complementing the *ΔrpoD4* mutation restores the WT period in *sigC* expression (median ≈ 24.43 h, p = 0.4) (**Fig. S4C**), further giving us confidence that the *rpoD4* mutation causes a reduction in clock period.

We then confirmed that overexpression of RpoD4 causes an increase in clock period, as observed in bulk luciferase experiments by Nair *et al*. (Nair *et al*, 2002). We placed the *rpoD4* gene under the transcriptional control of the IPTG-inducible P*_trc_* promoter (**Fig. 3D**) and assayed the effects of RpoD4 overexpression on P*_kaiBC_*-EYFP-fsLVA expression under constant light conditions (c.a. 15 μE m^-2^ s^-1^). As in previously described period analysis, we extracted mean P*_kaiBC_*-EYFP expression traces from time-lapse movies of WT, non-induced P*_trc_*-*rpoD4* and 100 μM IPTG induced P*_trc_*-*rpoD4* (abbreviated as *rpoD4OX* in figures) reporter strains using coarse colony-level analysis (**Fig. 3C**). We then detrended and inputted the data into BioDare2 for period analysis (FFT-NLLS method, period window constrained to 22-28 h). The period of P*_kaiBC_*-EYFP oscillations was increased by about 1 h (p = 2.9e-13, Wilcoxon rank-sum test) in cells overexpressing *rpoD4* (median = 25.8 h) when compared to cells with non-induced P*_trc_*-*rpoD4* (median = 24.8 h) (**Fig. 3D**).

### RpoD4 levels modulate cell size

Given the pulsatile expression dynamics of RpoD4 during cell division, we speculated deletion or overexpression of RpoD4 might cause cell division and/or cell size phenotypes. *S. elongatus* is rod-shaped and grows in volume by increasing pole-to-pole length, so the cell length is a good proxy of cell size and the elongation rate is correlated with cell growth rate (Martins *et al*, 2018). To test whether *rpoD4* deletion caused a cell size phenotype, and whether this phenotype was dependent on the circadian clock or not, we examined WT, *ΔrpoD4*, *ΔkaiBC* and *ΔkaiBCΔrpoD4* strains carrying the P*_rpoD4_*-EYFP-LVA reporter and grown on agarose pads under constant light (c.a. 18 μE m^-2^ s^-1^). Under this condition, the median cell cycle durations of the four strains are almost identical (ranging from 14.25 to 15 h), and deletion of the *kaiBC* operon abolished the bimodal distribution of cell cycle durations (**Fig. S5A**). We found that deletion of *rpoD4* caused a small but statistically significant reduction in birth, division and added length in both WT and *ΔkaiBC* strains (**Fig. 4A-C**). This suggests that deletion of *rpoD4* causes a cell size reduction phenotype, which is not dependent on the circadian clock. Next, we analysed the dependency of birth and division length on time of day at birth and division, respectively, in those four strains. Birth and division lengths oscillate throughout the day, but deletion of *rpoD4* reduced cell length across all times of day (**Fig. S5B&C**). In a *ΔkaiBC* background, deletion of *rpoD4* again reduced birth and division length across the day (**Fig. S5D&E**). To confirm this effect of *rpoD4* deletion on cell size, we performed a genetic complementation of the *rpoD4*-deletion mutant (denoted *ΔrpoD4-rpoD4*), which restores the WT size phenotype (**Fig. 4D**). We also confirmed that a similar cell size reduction could be observed in cells grown in liquid culture rather than on agarose pads (**Fig. S5F**).

**Figure 4.**
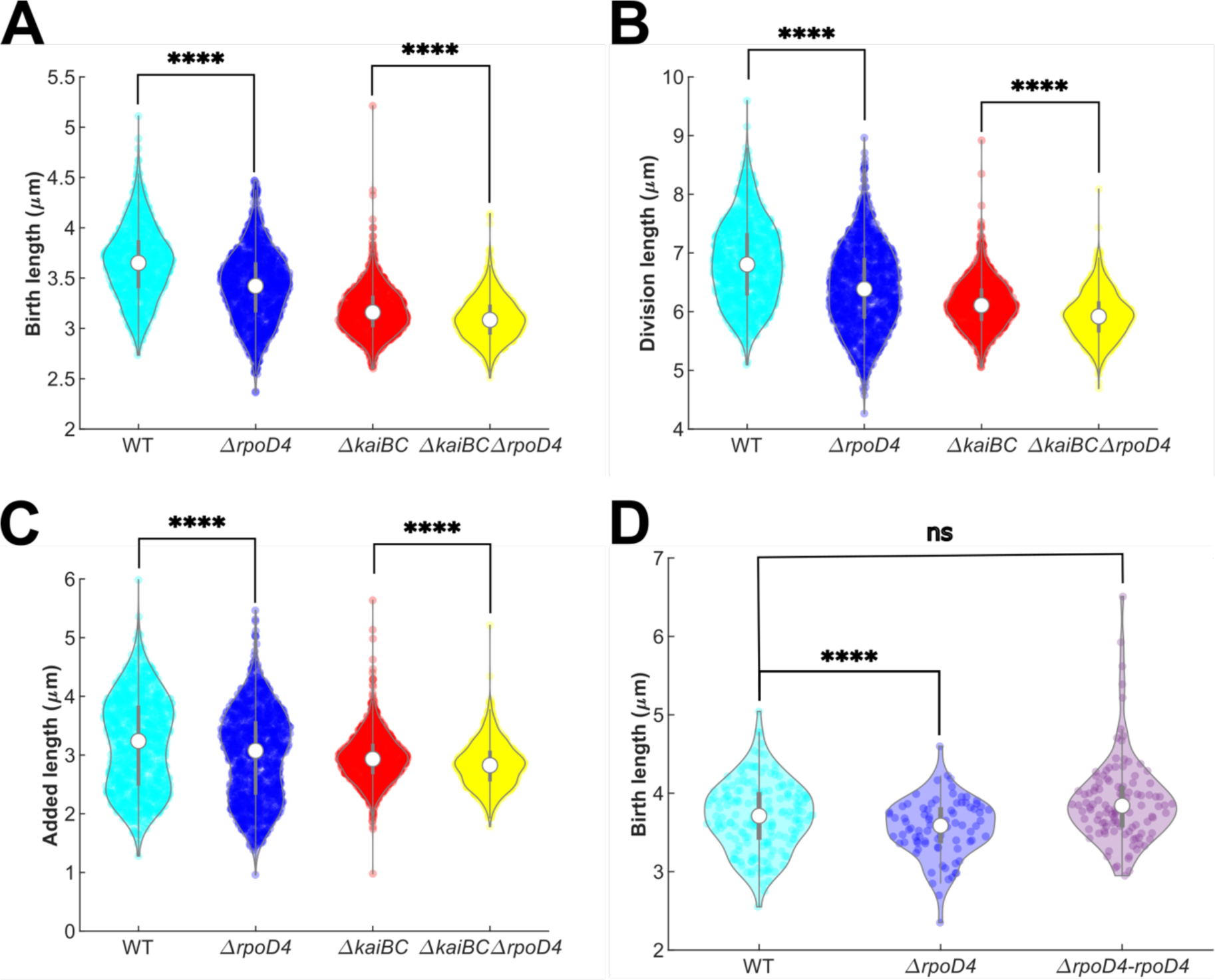
Deletion of *rpoD4* reduces cell size. Violin plots of (**A**) birth length, (**B**) division length, (**C**) added length of WT (cyan), *ΔrpoD4* (blue), *ΔkaiBC* (red) and *ΔrpoD4ΔkaiBC* (yellow) strains carrying P*_rpoD4_*-EYFP-LVA reporters grown on agarose pads under constant light (c.a. 18 μE m^-2^ s^-1^). For each plot, the central mark (white circle) indicates the median, and the top and bottom of the grey bar indicate the 25th and 75th percentiles, respectively. 1140 WT, 1310 *ΔrpoD4*, 1350 *ΔkaiBC*, and 1379 *ΔrpoD4ΔkaiBC* cells from two independent experiments were collected for analysis. (**D**) Violin plots of birth length of WT, *ΔrpoD4* and *ΔrpoD4-rpoD4* (purple) strains without any reporters grown under constant higher light (c.a. 45 μE m^-2^ s^-1^). 173 WT, 80 *ΔrpoD4* and 110 *ΔrpoD4-rpoD4* cells from one experiment were collected for analysis. ****: p < 0.0001, ns: not significant.

Because we observed *rpoD4*-deletion reduced cell size for cells grown on agarose pads and liquid cultures, we then asked whether overexpression of *rpoD4* has the opposite effect, leading to an increase in cell size. To answer this question, we examined the cell size distributions of the data shown in **Fig. 3** for WT and P*_trc_*-*rpoD4* strains. P*_trc_*-*rpoD4* strain was induced with 100 μM of IPTG to overexpress *rpoD4* throughout the experiment, and we refer to this condition as *rpoD4*OX. Under this condition, the cell cycle duration and elongation rate were close between WT and *rpoD4*OX (**Fig. 5A&B**). However, overexpression of *rpoD4* increased the birth, division and added length by 40%, 49% and 58%, respectively, relative to WT (**Fig. 5C-E**, **Fig. S6**). Because of the nature of exponential growth, *rpoD4OX* cells can maintain larger mean birth and division lengths relative to WT, while maintaining similar cell cycle duration and elongation rate.

**Figure 5.**
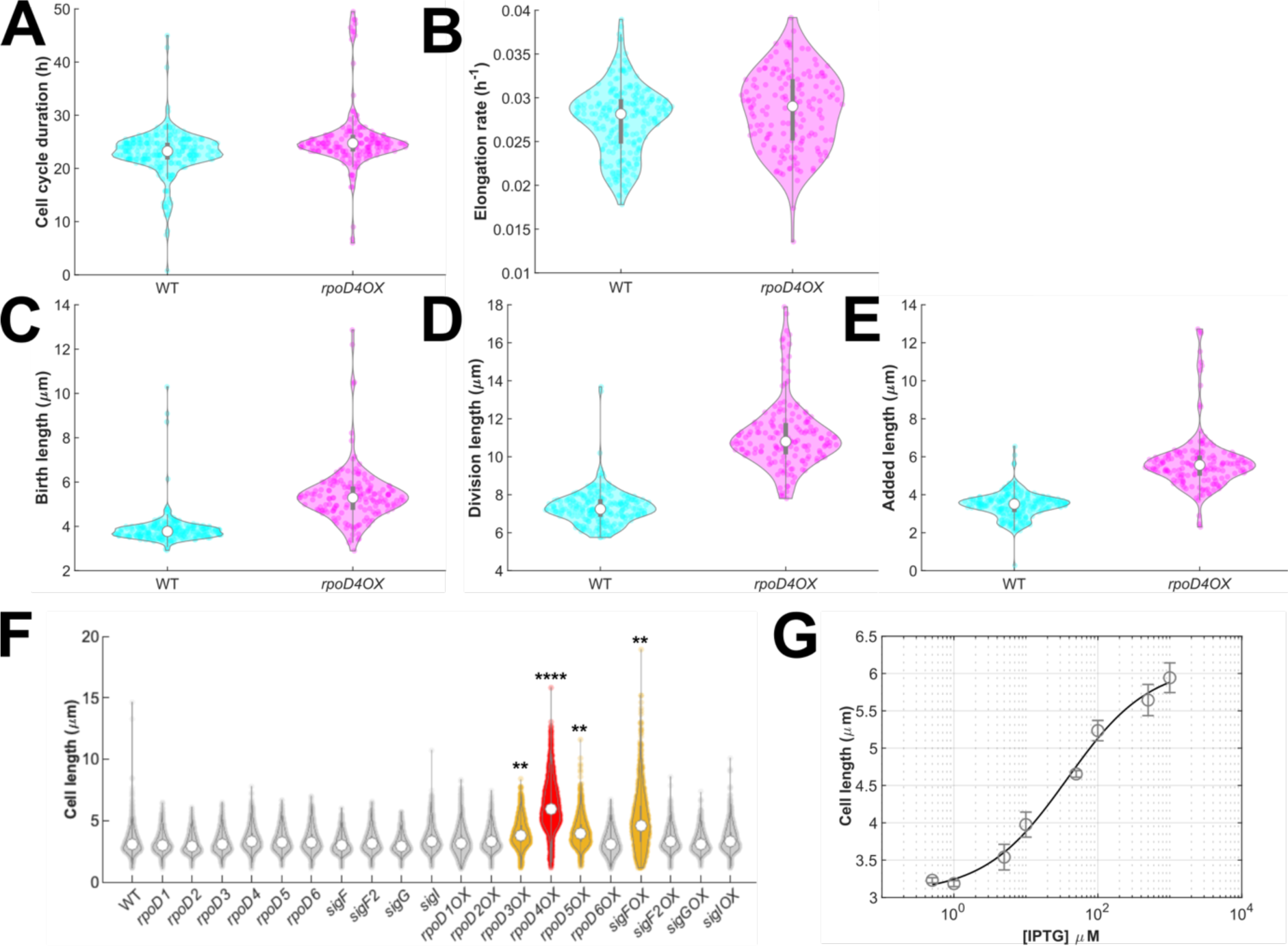
Overexpression of *rpoD4* increases cell size. Violin plots of (**A**) doubling time, (**B**) elongation rate, (**C**) birth length, (**D**) division length, and (**E**) added length of WT (blue) and 100 μM IPTG induced P*_trc_*-*rpoD4* (magenta, abbreviated as *rpoD4OX*) strains carrying P*_kaiBC_*-EYFP-fsLVA reporter and grown on agarose pads under constant light (c.a. 15 μE m^-2^ s^-1^). 239 WT and 161 *rpoD4OX* cells from one experiment were collected for analysis. (**F**) Cell length of WT and inducible strains for ten different sigma factors (P*_trc_*-sigma) incubated with or without 1 mM IPTG for 3 days under constant light (20-25 μE m^-2^ s^-1^) in liquid cultures. The strains incubated with and without IPTG are named as *sigma* and *sigmaOX*, respectively. Data collected from three independent experiments are shown as violin plots with the central mark indicating the median. Strains with significantly different cell size compared to WT are highlighted with different colours (**: p < 0.01 (yellow), ****: p<0.0001 (red)). (**G**) Cell length of P*_trc_*-*rpoD4* cells incubated with increasing concentration of IPTG for three days. Data shown as mean ± SEM of three independent experiments. A Hill equation is fitted to the data with EC_50_ = 37 μM and Hill coefficient = 0.758.

Although we observed overexpression of *rpoD4* increased cell size, it was possible that the cell size difference might not be specific to *rpoD4*, but could be caused by the overexpression of any alternative sigma factor. Therefore, we did control experiments in which we generated overexpression strains for all known sigma factors in *S. elongatus* (P*_trc_*-sigma). We grew WT and P*_trc_*-sigma strains with or without 1 mM IPTG in batch cultures from an initial OD_750_ of 0.05. After three days of cultivation under constant ambient light (20-25 µE m^-2^ s^-1^), cells were sampled for microscope snapshots and size analysis. As shown in **Fig. 5F**, overexpression of only 4 of the 10 alternative sigma factors caused a significant cell size increase, and the *rpoD4OX* strain had the most significant increase amongst all sigma factors. The mean cell length of *rpoD4*OX strain (6.17 μm) is almost doubled compared to WT (3.22 μm). We note that in these experiments, growth conditions in liquid culture (**Fig. 5F&G**) seem to amplify the cell size difference relative to growth in agarose pads (**Fig. 5A-E**). We also checked the OD_750_ of all strains, which could affect the cell size when they are sampled. Most strains grew to mid-exponential phase (OD_750_ = 0.4-0.8), with only the *sigF*OX strain struggling to grow, reaching an average OD_750_ = 0.2 after 3 days (**Fig. S7A**). By checking the snapshots of the sampled strains, we found there are many small as well as filamentous *sigF*OX cells, which is a possible indication of stress. In contrast, the *rpoD4*OX cells are longer but have a more uniform distribution (**Fig. S7B**). Taken together, the data suggests overexpression of *rpoD4* results in the most significant cell size increase among all sigma factors in *S. elongatus* and does not cause a growth defect.

Finally, we checked if there is a quantitative relationship between *rpoD4* expression level and cell size. We grew the P*_trc_*-*rpoD4* strain with increasing concentration of IPTG in batch cultures under constant ambient light (20-25 µE m^-2^ s^-1^) for three days. The cells were then sampled for snapshots and size analysis (**Fig. S8**). EC_50_ = 37 μM and Hill coefficient = 0.758 (**Fig. 5H**). The results thus suggest that there is a dose-response relationship between the *rpoD4* expression level and cell size.

## Discussion

Using fluorescent protein reporters and quantitative single-cell microscopy, we have shown that the expression of the group 2 sigma factor *rpoD4* in *S. elongatus* pulses at cell division (**Fig. 1**), with *rpoD4* reporter pulse amplitude and pulse width modulated by the circadian clock. Pulses in *rpoD4* expression have, on average, higher amplitude in cell division events occurring around subjective dusk, but are wider at subjective dawn (**Fig. 2 A&C**). In contrast, pulses in clock-deletion cells (*ΔkaiBC*) have similar amplitudes and widths throughout the day (**Fig. 2B&D**). We verified that these dynamics were not an artefact of our transcriptional reporter and observed the same behaviour in cells carrying a protein fusion reporter instead (RpoD4-EYFP, **Fig. S1**).

We found that RpoD4 feeds back to the clock, as manipulating *rpoD4* levels alters the period of two transcriptional reporters of the circadian clock (P*_kaiBC_*-EYFP and P*_sigC_*-EYFP). Constitutive overexpression of *rpoD4* leads to lengthening of clock period by ca. 1.1 h (**Fig. 3C&D** and **Fig. S4**, with an extra copy of *rpoD4* driven by the *trc* promoter with 100 μM IPTG induction), as similarly reported by *Nair et al.* (Nair *et al*, 2002). However, we also observed a slight, but statistically significant, shortening of the clock period in *rpoD4*-deletion cells (**Fig. 3AB** and **Fig. S4**), suggesting an inversely proportional relation between levels of RpoD4 and clock period. While expression of endogenous *rpoD4* peaks in a window of only a few hours around cytokinesis, and so its cycle of expression is synchronous with the cell division cycle, the clock has a natural period of about 24 h. In our experiments, cell cycle durations can be as low as 14.25 h on average (**Fig. S5A**, 18 μE m^-2^ h^-1^), and so it is intriguing to speculate whether these two oscillators, the clock and the cell division cycle, can cross-talk via RpoD4, at least indirectly. Clock regulation of the timing of cell division, and thus of the cell cycle, is well established (Dong *et al*, 2010; Mori *et al*, 1996; Martins *et al*, 2018; Yang *et al*, 2010), but it is believed the clock itself is largely insulated from cell division-induced fluctuations (Mori & Johnson, 2001; Paijmans *et al*, 2016; Yang *et al*, 2010). In future, it will be interesting to test whether upregulated, but temporally restricted to the window around cytokinesis, expression of *rpoD4* causes similar changes in clock period. A better understanding of sigma factor regulons and the cross-regulatory interactions between alternative sigma factors (Fleming & O’Shea, 2018) will also shed light on those questions.

Revealing the regulatory mechanisms of sigma factor dynamics in *S. elongatus* can provide new methods for controlling the oscillatory dynamics of the clock and allow the design of new genetic circuits for synthetic biology applications. For instance, the *rpoD4* promoter could be used as a building block to drive the expression of genes of interest specifically at cell division in *S. elongatus.* Future work should focus on this question, aiming to uncover the specific mechanisms of *rpoD4* activation and its downstream regulons. Furthermore, if we could elucidate these, RpoD4 and the *rpoD4* promoter could be used for orthogonal gene expression in other bacteria. This could help optimise the production of valuable (bio)chemicals in better studied synthetic biology chassis, such as *Escherichia coli* (Bervoets *et al*, 2018).

Since *rpoD4* expression coincides with cell division and the timing of cell division affects cell size in cyanobacteria (Martins *et al*, 2018), we asked whether *rpoD4* affects cell size. *rpoD4* deletion causes a small reduction in average cell size (**Fig. 4**), while overexpression leads to larger cell sizes (**Fig. 5C-E**). We show that this dependency of cell size on RpoD4 levels follows a clear dose-response relation (**Fig. 5G**). While revealing all possible pleiotropic effects of manipulating RpoD4 levels in *S. elongatus* is outside the scope of our study, this dose-response relation offers an alternative form of experimentally manipulating cell size, with minimal impact on growth (**Fig. 5A&B**). This could lead to further insight in our understanding of the regulation of chromosome and protein copy numbers (Zheng & O’Shea, 2017; Jun & Rust, 2017), as well as in other open questions in cell physiology.

Studying the clock-sigma factor coupling in *S. elongatus* might also reveal general design principles of clock coupling that can be applied to other systems such as plants and mammals. One design principle of clock coupling revealed by our work is that sigma factors can serve as hubs to integrate information from the clock and other cellular processes (e.g., cell division, as in the case of RpoD4). In support of this idea, one of the sigma factors in plants, SIG5, has been shown to integrate light and circadian clock signals to regulate gene expression in chloroplasts (Belbin *et al*, 2017; Noordally *et al*, 2013). It would be interesting to investigate whether other plant sigma factors, which have evolutionary connections with cyanobacteria sigma factors (Cuitun-Coronado & Dodd, 2020), could also interact with the circadian clock to regulate other cellular processes in plants.

Taken together, the results in our work reveal cell-division correlated pulsing dynamics of the alternative sigma factor RpoD4 in cyanobacteria, and shed light on links between RpoD4, the clock and the cell cycle. This highlights the value of single-cell analysis, as these interconnections are obscured by bulk averaged approaches. Our findings pave the way for better understanding of how the circadian clock regulates and receive inputs from a wide range of physiological processes in cyanobacteria and other organisms.

## Materials and methods

### Bacterial strains

*Synechococcus elongatus* PCC 7942 (ATCC® 33912) (*S. elongatus*) was used as the wild-type (WT) strain for genetic manipulations. Chemically competent *Escherichia coli* (*E. coli*) *DH*5α and HST08 (Stellar Competent Cells, TaKaRa Bio Inc., Kusatsu, Shiga, Japan) were used as the host strains for molecular cloning.

### Media and growth conditions

For the propagation of plasmids, *E. coli* was grown in Luria-Bertani (LB) medium (Bertani, 1951) with appropriate antibiotics for 18 hours at 37 °C with constant rotation at 220 rpm. *E. coli* plate cultures for obtaining single colonies was carried out for 18 hours at 37 °C, on 20 mL LB-agar plates containing appropriate antibiotics. Antibiotic working concentrations in LB media were as follows: 100 µg mL^-1^ ampicillin (Amp), 50 µg mL^-1^ gentamicin (Gent), 50 µg mL^-1^ kanamycin (Kan), 100 µg mL^-1^ spectinomycin (Spt), and 100 µg mL^-1^ chloramphenicol (Chl).

Modified BG-11 (BG-11 M) liquid medium (Mackey *et al*, 2007) with appropriate antibiotics was used for cultivating *S. elongatus*. Liquid cultures were grown at 30 °C under constant light of photosynthetically active radiation (PAR) = 20-25 µE m^-2^ s^-1^ and rotation at 120 rpm. *S. elongatus* plate cultures for obtaining and patching single colonies were carried out on 40 mL BG-11 M solid medium (Mackey *et al*, 2007) without Na_2_SO_3_, and the plate cultures were maintained under a light intensity of ∼100 µE m^-2^ s^-1^. Antibiotic working concentrations in BG-11 M media were as follows: 2 µg mL^-1^ Spt, 2 µg mL^-1^ streptomycin (Str), 2 µg mL^-1^ Gent, 5 µg mL^-1^ Kan, and 7.5 µg mL^-^ _1_ Chl.

### DNA manipulation and strain construction

Molecular cloning was done by standard restriction enzyme digestion cloning, Gibson assembly (Gibson *et al*, 2009) or In-Fusion® cloning (Irwin *et al*, 2012) (TaKaRa Bio Inc., Kusatsu, Shiga, Japan). *S. elongatus* transformation was achieved by homologous recombination following the protocol by Mackey *et al*. (Mackey *et al*, 2007) with modifications. Briefly, a 3 mL culture at log phase was pelleted by centrifuging at 4000 rpm for 5 mins. The pelleted cells were washed with 3 mL of 10 mM NaCl and pelleted again. The washed cells were resuspended in 300 μL of fresh BG-11M medium and mixed with 1 μg of the plasmid. Complete allele replacement was confirmed by polymerase chain reaction (PCR) and gel electrophoresis of the genomic DNA extracted from the transformed strains patched on solid media or grown in liquid cultures. Genomic DNA extraction was done by either boiling cells in 25 μL of 20 mM NaOH at 95 °C for 15 min or by using the Direct DNA Extraction Kit (Bacteria) (Norgen Biotek, Thorold, ON, Canada) following the manufacturer’s manual.

All transcriptional reporters were constructed using 1000 bp (except for the shorter promoter length used in **Fig. S3**) upstream of the start codon of the gene of interest driving the expression of fluorescent proteins as described in the text. For the P*_rpoD4_*-RpoD4-EYFP protein fusion reporter, we fused P*_rpoD4_*-*rpoD4* (*rpoD4* ORF with 1000 bp upstream of it as the *rpoD4* promoter) to *eyfp* via a flexible linker (GSGSGSG). Gene deletion was achieved by disruption of the endogenous gene with a gentamicin-resistance cassette (a gift from Prof. Erin O’Shea), and kanamycin-resistance cassette (from pUC19) through homologous recombination. The plasmids and *S. elongatus* strains used in this study are listed in **Table 1** and **Table 2**.

**Table 1.**
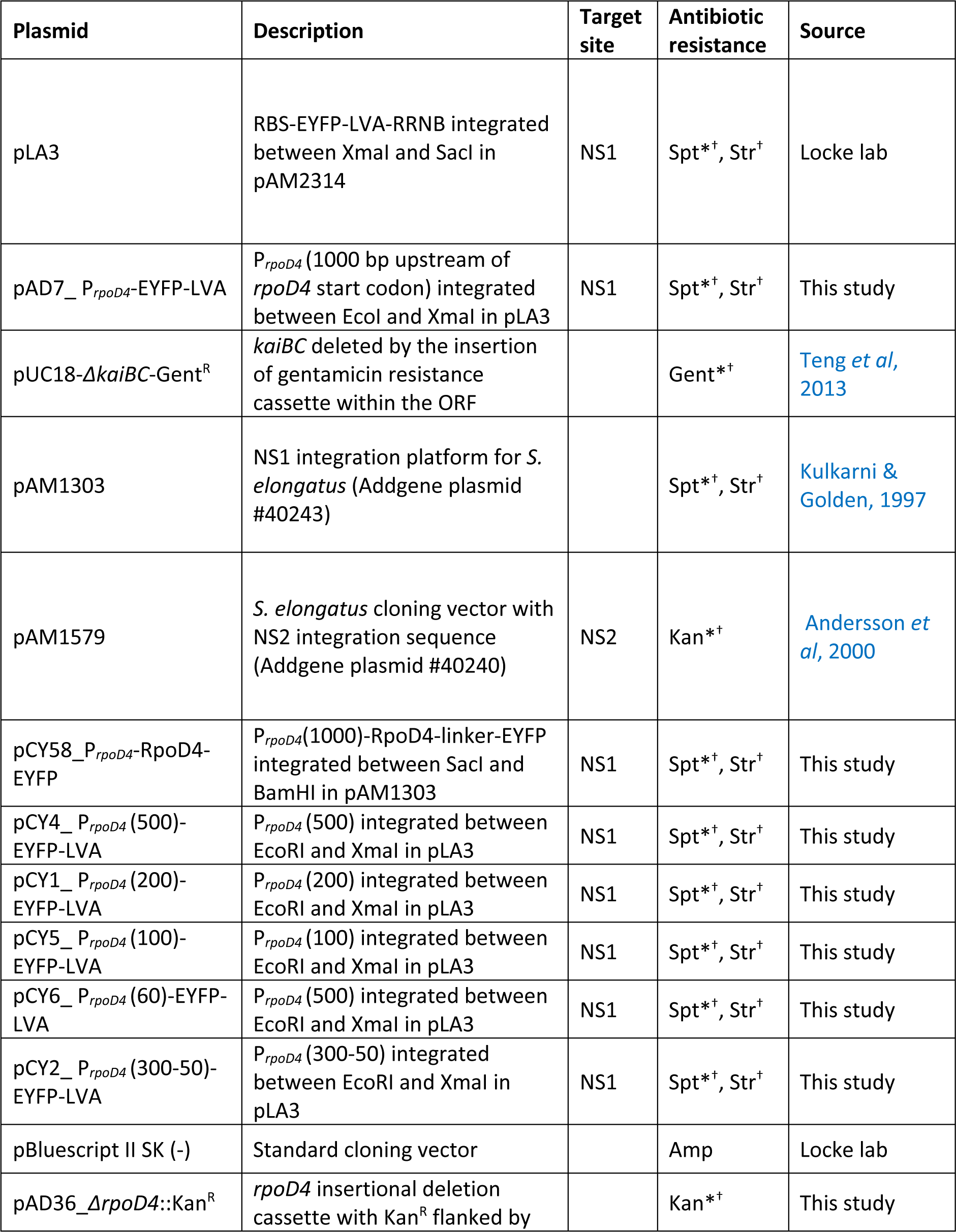
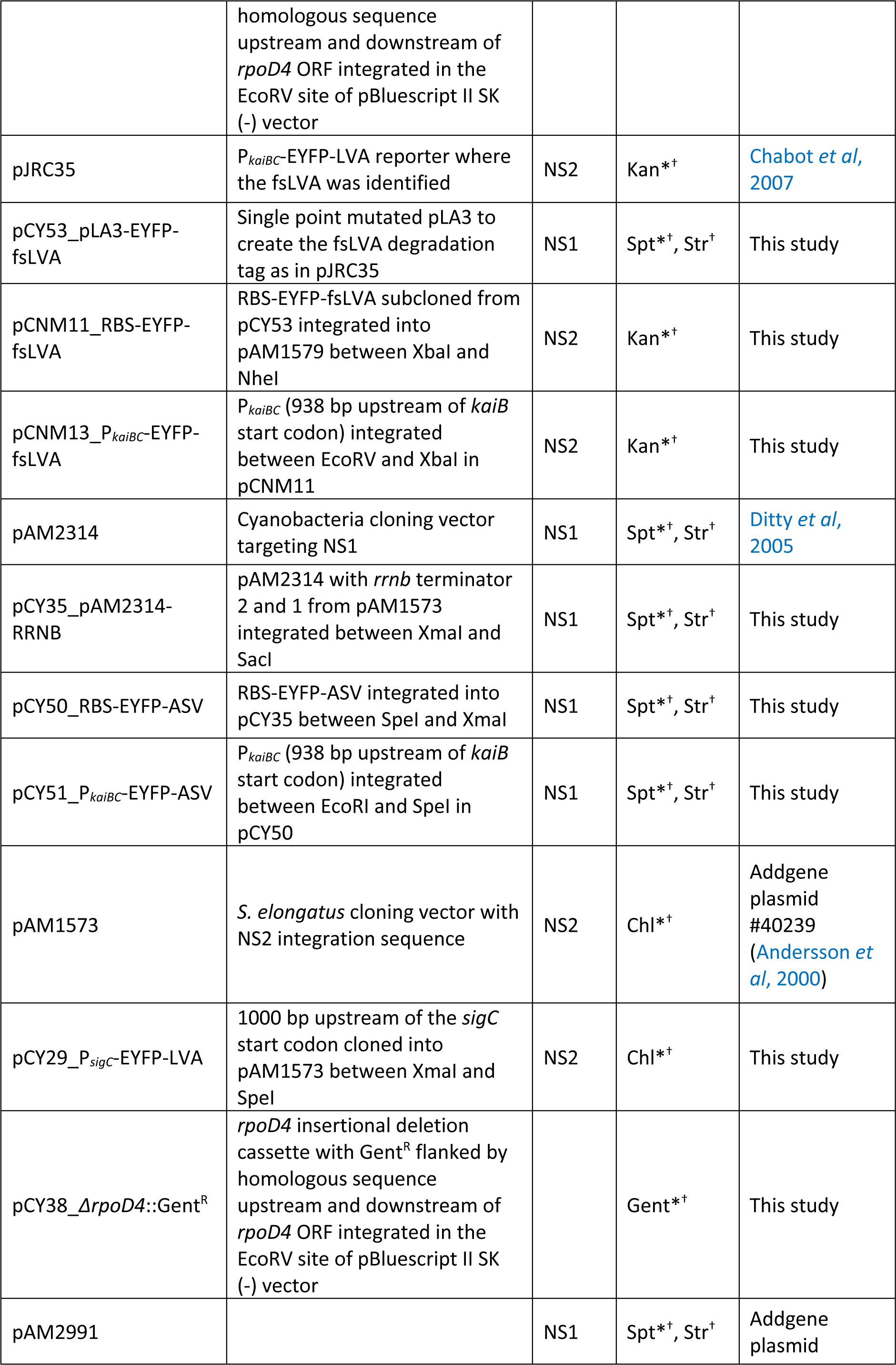
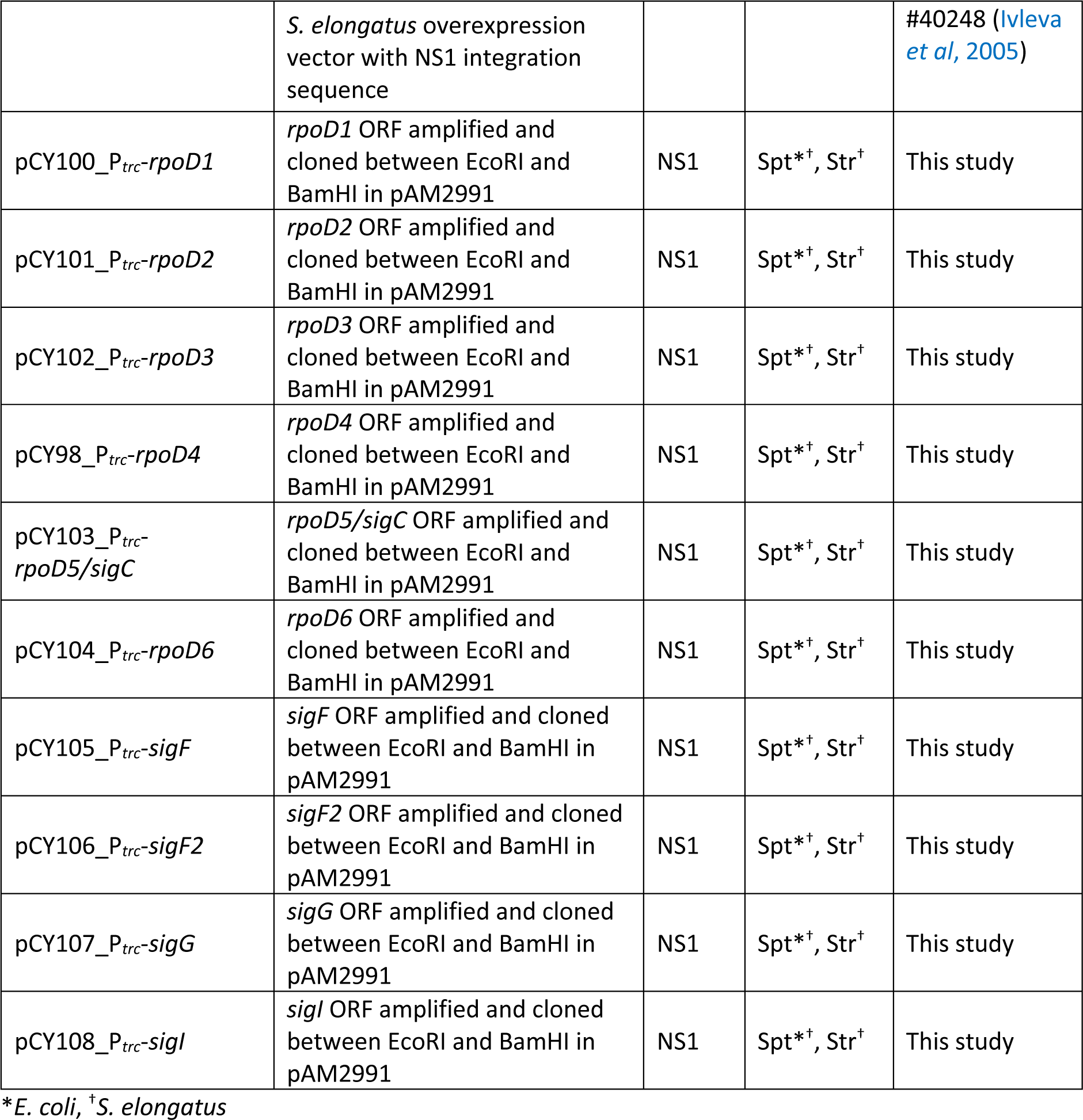
Plasmids used in this study.

**Table 2.**
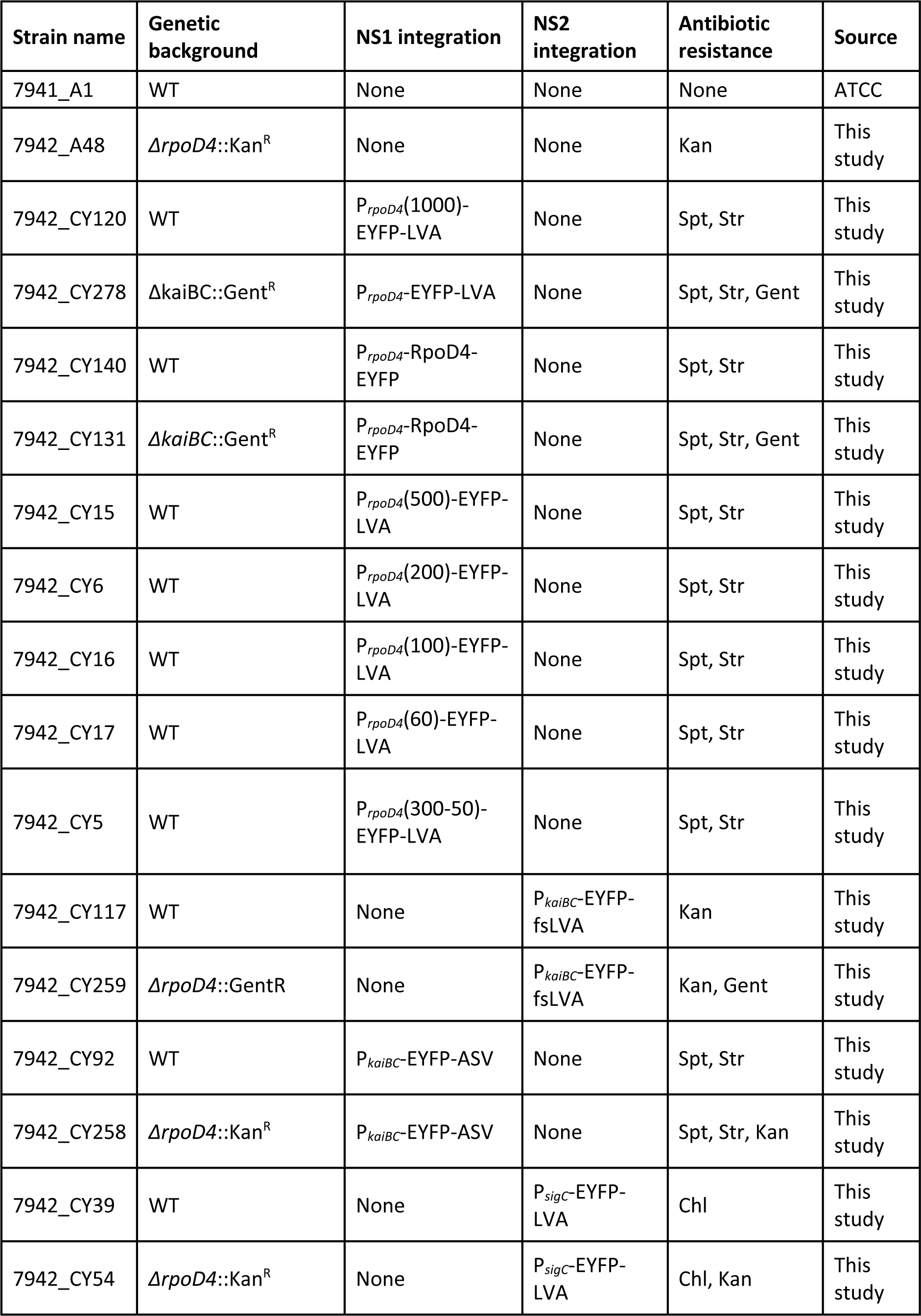
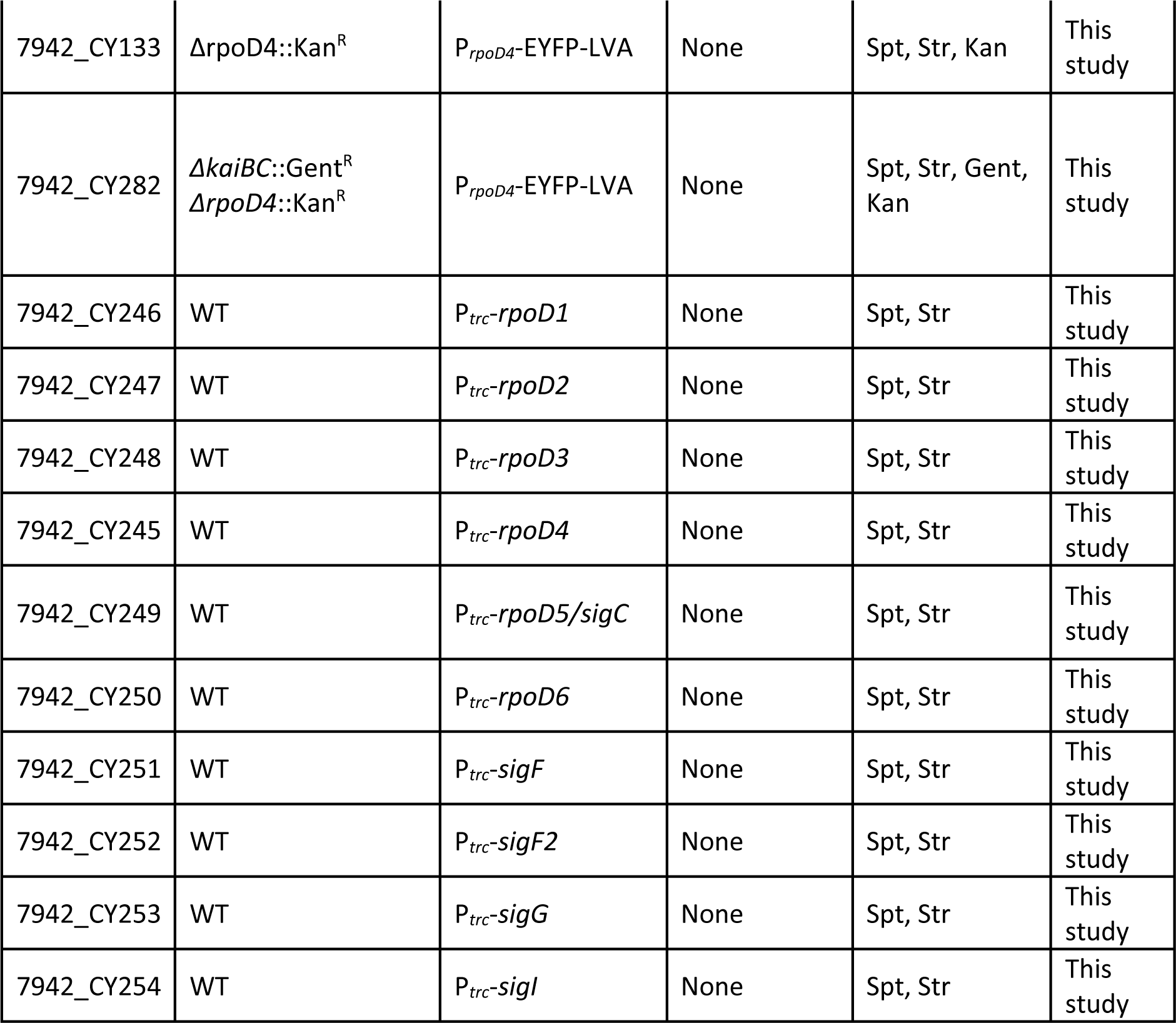
*S. elongatus* strains used in this study.

| Strain name | Genetic background | NS1 integration | NS2 integration | Antibiotic resistance | Source |
| --- | --- | --- | --- | --- | --- |
| 7941_A1 | WT | None | None | None | ATCC |
| 7942_A48 | $\Delta rpoD4::Kan^R$ | None | None | Kan | This study |
| 7942_CY120 | WT | P<sub>rpoD4</sub>(1000)-EYFP-LVA | None | Spt, Str | This study |
| 7942_CY278 | $\Delta kaiBC::Gent^R$ | P<sub>rpoD4</sub>-EYFP-LVA | None | Spt, Str, Gent | This study |
| 7942_CY140 | WT | P<sub>rpoD4</sub>-RpoD4-EYFP | None | Spt, Str | This study |
| 7942_CY131 | $\Delta kaiBC::Gent^R$ | P<sub>rpoD4</sub>-RpoD4-EYFP | None | Spt, Str, Gent | This study |
| 7942_CY15 | WT | P<sub>rpoD4</sub>(500)-EYFP-LVA | None | Spt, Str | This study |
| 7942_CY6 | WT | P<sub>rpoD4</sub>(200)-EYFP-LVA | None | Spt, Str | This study |
| 7942_CY16 | WT | P<sub>rpoD4</sub>(100)-EYFP-LVA | None | Spt, Str | This study |
| 7942_CY17 | WT | P<sub>rpoD4</sub>(60)-EYFP-LVA | None | Spt, Str | This study |
| 7942_CY5 | WT | P<sub>rpoD4</sub>(300-50)-EYFP-LVA | None | Spt, Str | This study |
| 7942_CY117 | WT | None | P<sub>kaiBC</sub>-EYFP-fsLVA | Kan | This study |
| 7942_CY259 | $\Delta rpoD4::GentR$ | None | P<sub>kaiBC</sub>-EYFP-fsLVA | Kan, Gent | This study |
| 7942_CY92 | WT | P<sub>kaiBC</sub>-EYFP-ASV | None | Spt, Str | This study |
| 7942_CY258 | $\Delta rpoD4::Kan^R$ | P<sub>kaiBC</sub>-EYFP-ASV | None | Spt, Str, Kan | This study |
| 7942_CY39 | WT | None | P<sub>sigC</sub>-EYFP-LVA | Chl | This study |
| 7942_CY54 | $\Delta rpoD4::Kan^R$ | None | P<sub>sigC</sub>-EYFP-LVA | Chl, Kan | This study |
| 7942_CY133 | $\Delta rpoD4::Kan^R$ | $P_{rpoD4}$-EYFP-LVA | None | Spt, Str, Kan | This study |
| 7942_CY282 | $\Delta kaiBC::Gent^R$<br>$\Delta rpoD4::Kan^R$ | $P_{rpoD4}$-EYFP-LVA | None | Spt, Str, Gent, Kan | This study |
| 7942_CY246 | WT | $P_{trc}$-*rpoD1* | None | Spt, Str | This study |
| 7942_CY247 | WT | $P_{trc}$-*rpoD2* | None | Spt, Str | This study |
| 7942_CY248 | WT | $P_{trc}$-*rpoD3* | None | Spt, Str | This study |
| 7942_CY245 | WT | $P_{trc}$-*rpoD4* | None | Spt, Str | This study |
| 7942_CY249 | WT | $P_{trc}$-*rpoD5/sigC* | None | Spt, Str | This study |
| 7942_CY250 | WT | $P_{trc}$-*rpoD6* | None | Spt, Str | This study |
| 7942_CY251 | WT | $P_{trc}$-*sigF* | None | Spt, Str | This study |
| 7942_CY252 | WT | $P_{trc}$-*sigF2* | None | Spt, Str | This study |
| 7942_CY253 | WT | $P_{trc}$-*sigG* | None | Spt, Str | This study |
| 7942_CY254 | WT | $P_{trc}$-*sigI* | None | Spt, Str | This study |

### Microscopy and sample preparation

#### Microscope setup

The microscope setup followed the method described by Martins *et al*. (Martins *et al*, 2016) with modifications. A Nikon Ti-E inverted microscope equipped with the Nikon Perfect Focus System modules was used to acquire images of *S. elongatus*. The temperature around the sample was maintained at 30 °C by a Solent environmental chamber (Solent Scientific, Portsmouth, UK). Light illumination was provided by a custom-built circular cool white light LED array (Cairn Research, Faversham, UK) attached to the condenser lens. Epi-illumination was provided by a solid-state white light source (Lumencor, Oregon, USA). Filter sets for fluorescent channels include 41027 Calcium Crimson (EX: 580/20x, EM: 630/60m), 49014 ET-mKO/mOrange (EX: 530/30x, EM: 575/40m), 49003 ET-EYFP (EX: 500/20x, EM: 535/30m), 49002 ET-GFP (EX: 470/40x, EM: 525/50m) and 49001 ET-CFP (EX: 436/20x, EM: 480/40m) from Chroma Technology (Vermont, USA). Phase-contrast and fluorescent images were acquired by a CoolSNAP HQ2 camera or a Prime sCMOS camera (Teledyne Photometrics, Arizona, USA). Data acquisition was controlled by the software Metamorph (Molecular Devices, California, USA). All experiments used a Nikon Plan Apochromat 100x objective and Nikon type-A immersion oil.

#### Agarose pad setup

The setup was modified from the protocol of Young *et al*. (Young *et al*, 2012). For time-lapse movies, 10 mL of cell culture in BG 11 M with a starting OD_750_ of 0.1 was entrained by at least two 12:12 h square LD cycles. Cells were then grown to an OD_750_ of 0.4 and diluted to an OD_750_ of 0.1. 1 μL of the diluted culture was spread evenly on 1.5% (w/v) BG-11 M agarose pads. The agarose pads were made from OmniPur® Low-Melting Agarose (Merk Millipore, CAS 9012-36-6). After drying, the agarose pads were placed face down into a two-chambered coverglass (Labtek Services, UK) and covered by 2.25 mL of melted BG-11 M agarose to prevent evaporation during the experiment. For all time-lapse microscopy experiments, samples were subject to a 12:12 h square LD cycle using the custom circular LED array before movie acquisition to synchronise their circadian clocks.

For snapshots, unless otherwise stated, cell cultures at exponential phase (OD_750_ = 0.4-0.8) were diluted to OD_750_ = 0.3. 2 μL of the diluted culture was spread evenly on 1.5% (w/v) BG-11 M agarose pads and transferred to Willco dishes (Willco Wells, Amsterdam) for image acquisition. Typically, 45 random positions were acquired automatically for each strain.

### Microscopy data analysis

#### Time-lapse movie analysis

For single-cell analysis of the agarose pad time-lapse movies, images were processed and analysed by a custom-made software in MATLAB R2014b and R2020a (The MathWorks, Massachusetts) adapted from Schnizcells 2.0 (Young *et al*, 2012), for segmentation and single-cell tracking.

The quantitative information of the micro-colonies was extracted from Schnitzcells-compiled data using code written in MATLAB R2020b (The MathWorks, Massachusetts). The data used in this study and how they were calculated are listed below.

**1) Mean fluorescence per cell:** Fluorescence quantification followed the protocol as described by Martins *et al*. (Martins *et al*, 2016). Briefly, mean fluorescence per cell, which can be used as an estimation of the gene expression level, is defined as the average pixel intensity within the segmented area of each cell. For movies of P*_kaiBC_* and P*_sigC_* reporters, all single-cell traces (i.e., mean fluorescence per cell) were smoothed by a moving average filter spanning five data points. Whenever the range of the filter window is not contained within the range of a cell cycle (i.e. when the filter is centred in one of the first two or in one of the last two data points of each individual cell cycle), single-cell traces are padded with data from the mother (on the left) and the average of the two daughters (on the right), provided data for these cells have been acquired.
**2) Cell length at birth and division:** the cell length is defined as the length of the major axis of the cell. The cell length at birth is the length of cell just after division (i.e., the first cell in a complete cell cycle). The cell length at division is the length of cell just before division (i.e., the last cell in a complete cell cycle).
**3) Added length:** defined as the increased length within a complete cell cycle and is calculated by division length minus birth length.
**4) Cell cycle duration:** defined as the timespan of a complete cell cycle from birth to division.
**5) Elongation rate:** defined as the extension rate of the major axis of the cell. It is calculated by extracting the exponent of an exponential curve fitted to the growth (in length) curve of a complete cell cycle.

#### Snapshot analysis

The snapshot analysis software was based on Schnitzcells 2.0 (Young *et al*, 2012) and adapted from Schwall *et al*. (Schwall *et al*, 2021). Briefly, the acquired image files were fed into the software and renamed. Cell segmentation was then carried out using the auto-fluorescence signal in Calcium Crimson channel images. Properties of the cells, including the major axis length, minor axis length, eccentricity, location of the centroid, mean fluorescence, and standard deviation of the fluorescence, were computed and organised into MATLAB structures as outputs.

Cell segmentation is not error-free without manual correction, but manual correction is time-consuming. To mitigate segmentation errors without doing extensive manual corrections, we developed a method to remove inaccurately segmented cells. First, we removed segmented objects whose centroid were more than 800 pixels away from the centre of the image, ensuring analysed cells were within the region of ≥80% epi-fluorescence illumination intensity. Next, we removed segmented objects whose minor axis length (width) was below 15 pixels (1 μm) and greater than 37.5 pixels (2.5 μm), because the width of *S. elongatus* grown in liquid cultures is normally within this range. After the two operations, the remaining segmented objects usually displayed consistent relationships between length and eccentricity (a non-negative real number that uniquely characterizes the shape of the cell; circular cells have eccentricity ∼ 0 and rod-shaped cells have eccentricity ∼ 1). We confirmed by visual inspection that most segmented objects that fell out of the selection criteria formed distinct clusters that did not overlap with the remaining cells.

### Quantitative analysis

#### Analysis of rpoD4 expression peaks

P*_rpoD4_*-EYFP expression profiles were extracted from all lineages in the dataset, then detrended by subtracting a second-order polynomial fit. The reason to detrend the data was to remove the effect of the systematic background fluorescence increase, which is caused by fluorescence bleed-through from neighbouring cells, and noticeable in the mean P*_rpoD4_*-EYFP expression trace of both WT and *ΔkaiBC* reporter strains (**Fig. S2A&B**). The increase in background fluorescence also occurs in a WT strain with no reporter and so is not due to a basal increase in *rpoD4* expression (**Fig. S9**). The *rpoD4* expression peaks were identified using the *findpeaks* function in MATLAB with minimum peak prominence ≥ 100 a.u., unless otherwise stated. The output from the *findpeaks* function include the location, height, width, and prominence of the peaks. Only unique peaks from all lineages were used for downstream analysis.

#### Circadian rhythmicity and period analysis

For circadian rhythmicity and period analysis, the time series were first detrended by subtracting a second-order polynomial fit, as we did for the *rpoD4* expression peaks. The detrended data were then exported to the BioDare2 suite (https://biodare2.ed.ac.uk/) (Zielinski *et al*, 2014). For rhythmicity analysis, we constrained the data window to 24-96 h, and the period window to 22-28 h. BD2 eJTK (Hutchison *et al*, 2015), (which provides accurate results for short data) and eJKT Classic (which works well for short data with an expected period of 22-26 h), were chosen for test method and analysis presets, respectively. For period analysis, we used the same data and period window as in the rhythmicity analysis, and selected FFT-NLLS (Plautz *et al*, 1997), which was developed for analysing data obtained in free-running conditions.

## Acknowledgements

We are grateful for plasmids provided by the O’Shea and Golden labs (see **Table 1**). Chris Micklem also aided in plasmid construction.

## Supplementary figures

**Figure S1.**
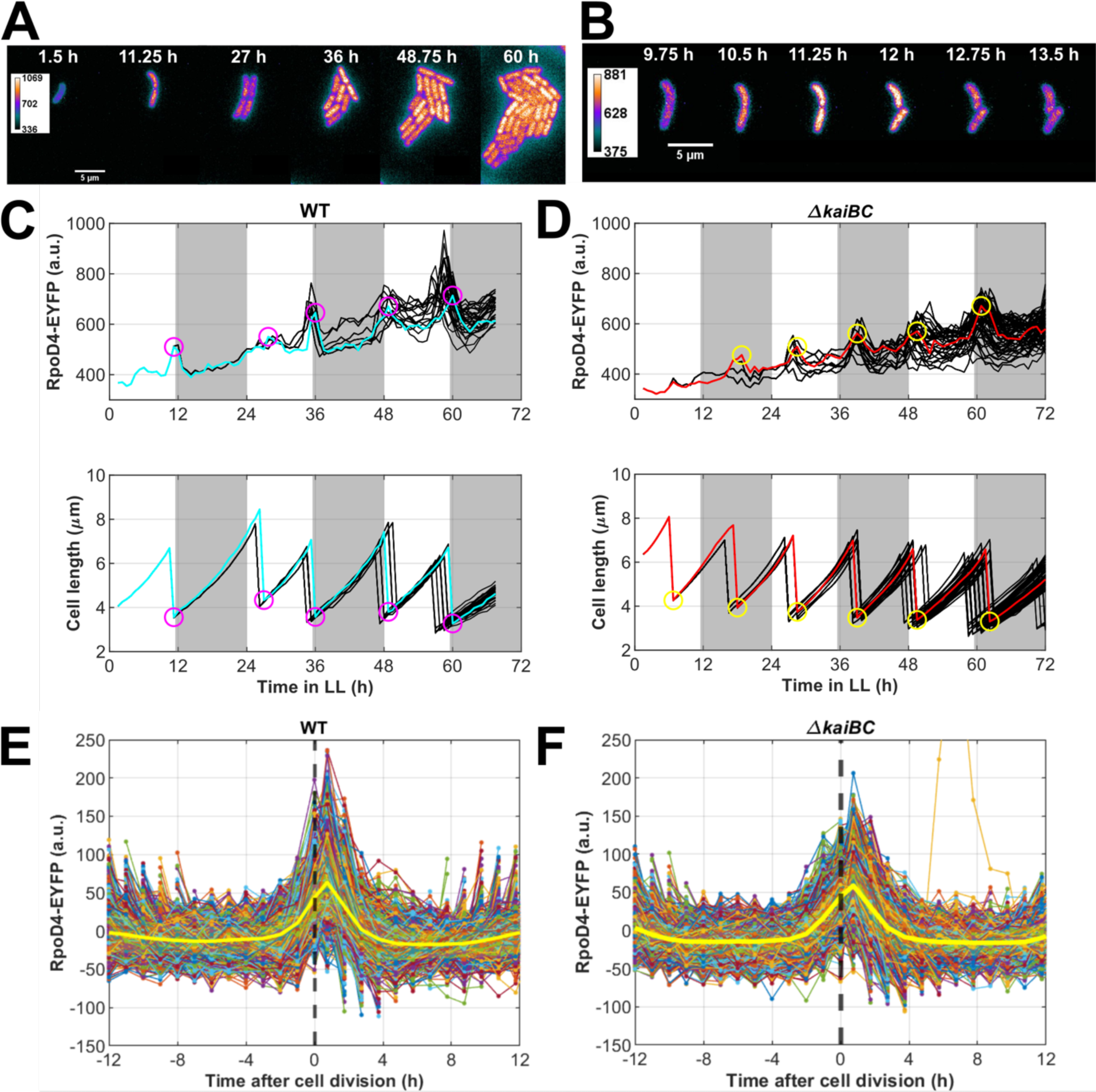
Protein levels of RpoD4-EYFP peak at cell division. (**A**) Montage of fluorescence microscopy images of WT P*_rpoD4_*-RpoD4-EYFP cells grown under constant light (c.a. 18 μE m^-2^ s^-1^). The heatmap indicates the YFP fluorescence intensity range. (**B**) Six consecutive frames from the same time-lapse movie as in (**A**) showing the expression of P*_rpoD4_*-RpoD4-EYFP pulses during cell division. (**C**) Profiles of P*_rpoD4_*-RpoD4-EYFP expression and cell length in WT or (**D**) *ΔkaiBC* background. Each line represents an individual cell or lineage, and the coloured lines are representative lineages. Note the peaks of P*_rpoD4_*-RpoD4-EYFP expression (prominence ≥ 40 a.u.) and the troughs of cell length profile are correlated as highlighted by coloured circles. White and grey shades represent subjective day and night, respectively. (**E**) Single-cell expression dynamics of P*_rpoD4_*-RpoD4-EYFP in WT and (**F**) *ΔkaiBC* background plotted against time centred at cell division (t = 0 h). Lines represent individual cells. The solid yellow line represents the mean. For both WT and *ΔkaiBC*, expression of P*_rpoD4_*-RpoD4-EYFP peaks just after cell division. 15 WT movies and 16 *ΔkaiBC* movies from 2 independent experiments were collected.

**Figure S2.**
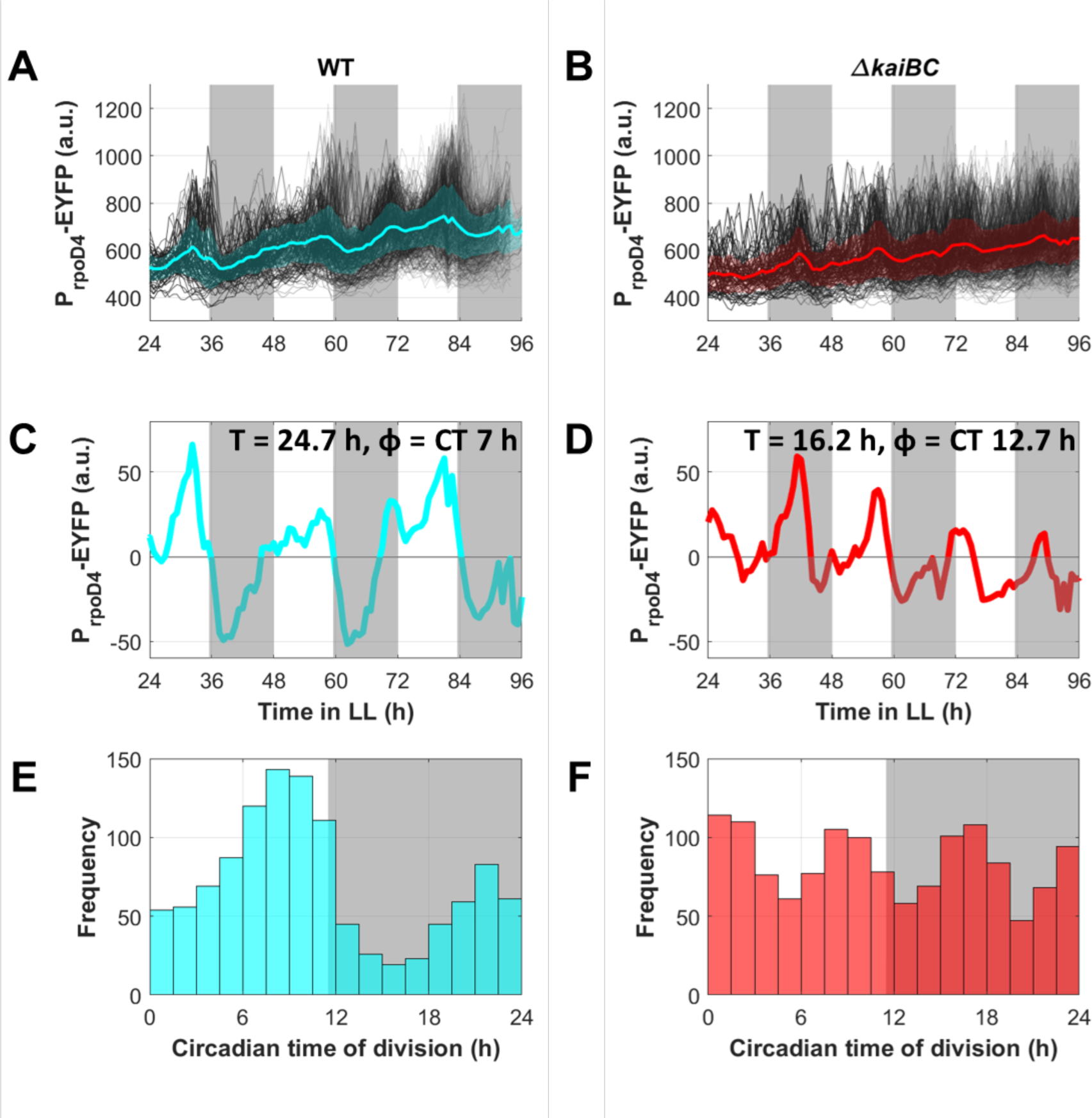
*rpoD4* expression displays circadian oscillations at the population level. (**A**) Time traces of P*rpoD4*-EYFP expression of individual lineages (black lines, 2373 cells from 19 movies in 2 independent experiments) and population average of WT P*rpoD4*-EYFP-LVA reporter strain (solid cyan line, with lighter cyan shades representing one standard deviation from the mean). (**B**) Time traces of P*rpoD4*-EYFP expression of individual lineages (black lines, 2775 cells from 18 movies in 2 independent experiments) and population average of P*rpoD4*-EYFP-LVA reporter in a *ΔkaiBC* background strain (solid red line, with lighter red shades representing one standard deviation from the mean). (**C**) Mean P*rpoD4*-EYFP expression of WT reporter strain after detrending. Rhythmicity analysis (test method: BD2 eJKT, analysis preset: eJKT Classic) confirmed the time series has circadian rhythmicity, with a period (T) of 24.67 h and an acrophase (φ) of CT 7.04 h. (**D**) Mean P*rpoD4*-EYFP expression of *ΔkaiBC* reporter strain after detrending. No circadian rhythm was detected by rhythmicity analysis, but period analysis suggests the time series has a period = 16.16 h. (**E**) Distribution of time of day at division of WT cells peaks towards the end of subjective day (peak at8.25 h). Data was collected from 1140 cell division events. (**F**) In contrast, division of *ΔkaiBC* cells is roughly uniform across time of day. Data was collected from 1350 cell division events.

**Figure S3.**
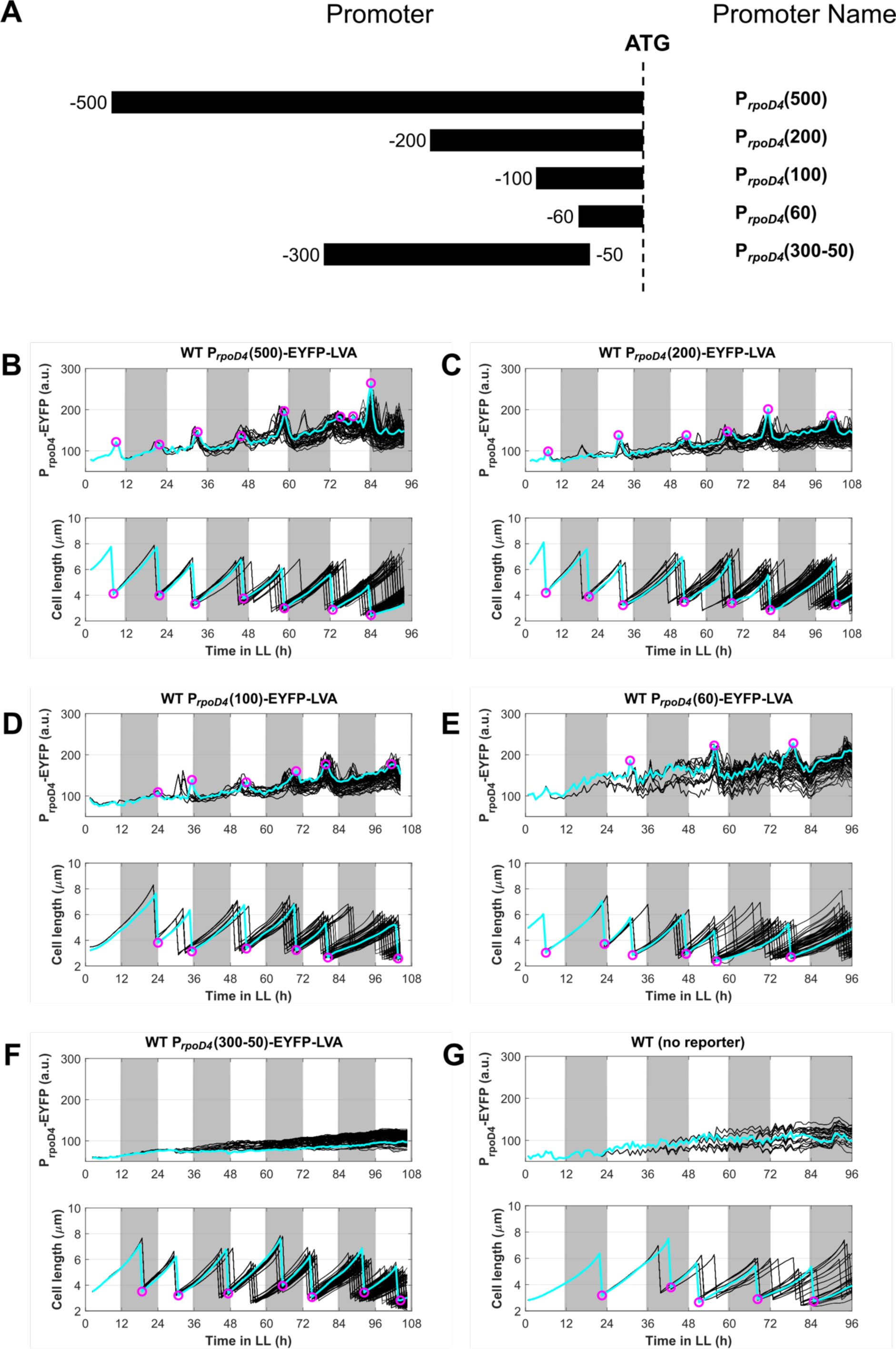
Cis-regulatory elements of *rpoD4* promoter locate in 100 bp upstream of the start codon. (**A**) Individual *rpoD4* promoter fragments with indicated endpoint (relative to the *rpoD4* start codon) were fused to EYFP-LVA to construct *rpoD4* transcriptional reporters. (**B-G**) WT with no reporters (G) and WT transformed with these reporters (B-F) were examined by time-lapse fluorescence microscopy on agarose pads under constant light conditions (c.a. 15 μE m-2 s-1). All apart from the truncated promoter P*_rpoD4_*(300-50) can drive EYFP-LVA expression above the WT background. One representative movie from n = 9 movies, shown for each strain.

**Figure S4.**
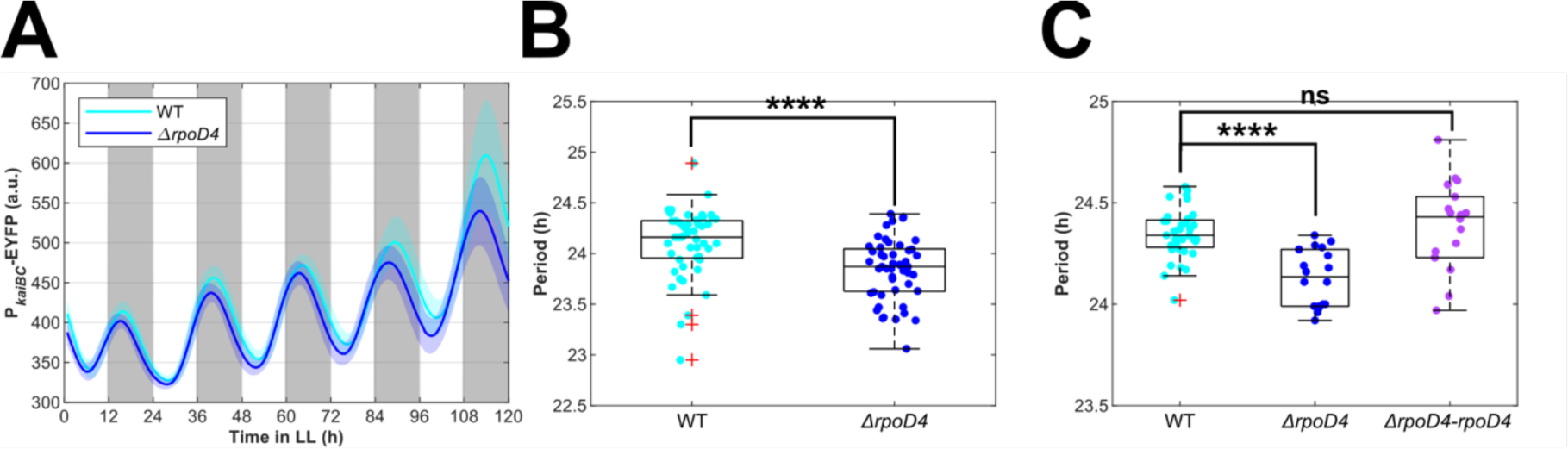
Deletion of *rpoD4* reduces the clock period. (**A**) Time trace of mean P*_kaiBC_*-EYFP-ASV expression of WT (cyan line, averaged from 49 movies) and *ΔrpoD4* (blue line, averaged from 49 movies) strains grown under constant high light conditions (c.a. 35 μE m^-2^ s^-1^). Data were collected from three independent experiments. (**B**) Box plot of the period of detrended mean P*_kaiBC_*-EYFP-ASV expression of WT (cyan dot, n = 49, median ≈ 24.16 h) and *ΔrpoD4* (blue dots, n = 49, median ≈ 23.87 h) movies. The difference is statistically significant (p = 2.8e-5). (**C**) Box plot of the period of detrended mean P*_sigC_*-EYFP-LVA expression of WT (cyan dot, n = 36, median ≈ 24.34 h), *ΔrpoD4* (blue dots, n = 18, median ≈ 24.14 h) and *ΔrpoD4-rpoD4* (purple dots, n = 18, median ≈ 24.43 h) movies. The difference between WT and *ΔrpoD4* is statistically significant (p = 4.23e-6), but the difference between WT and *ΔrpoD4-rpoD4* is not statistically significant (p = 0.4). Wilcoxon rank-sum test is used for all statistical tests in this figure. ****: p < 0.0001, ns: not significant.

**Figure S5.**
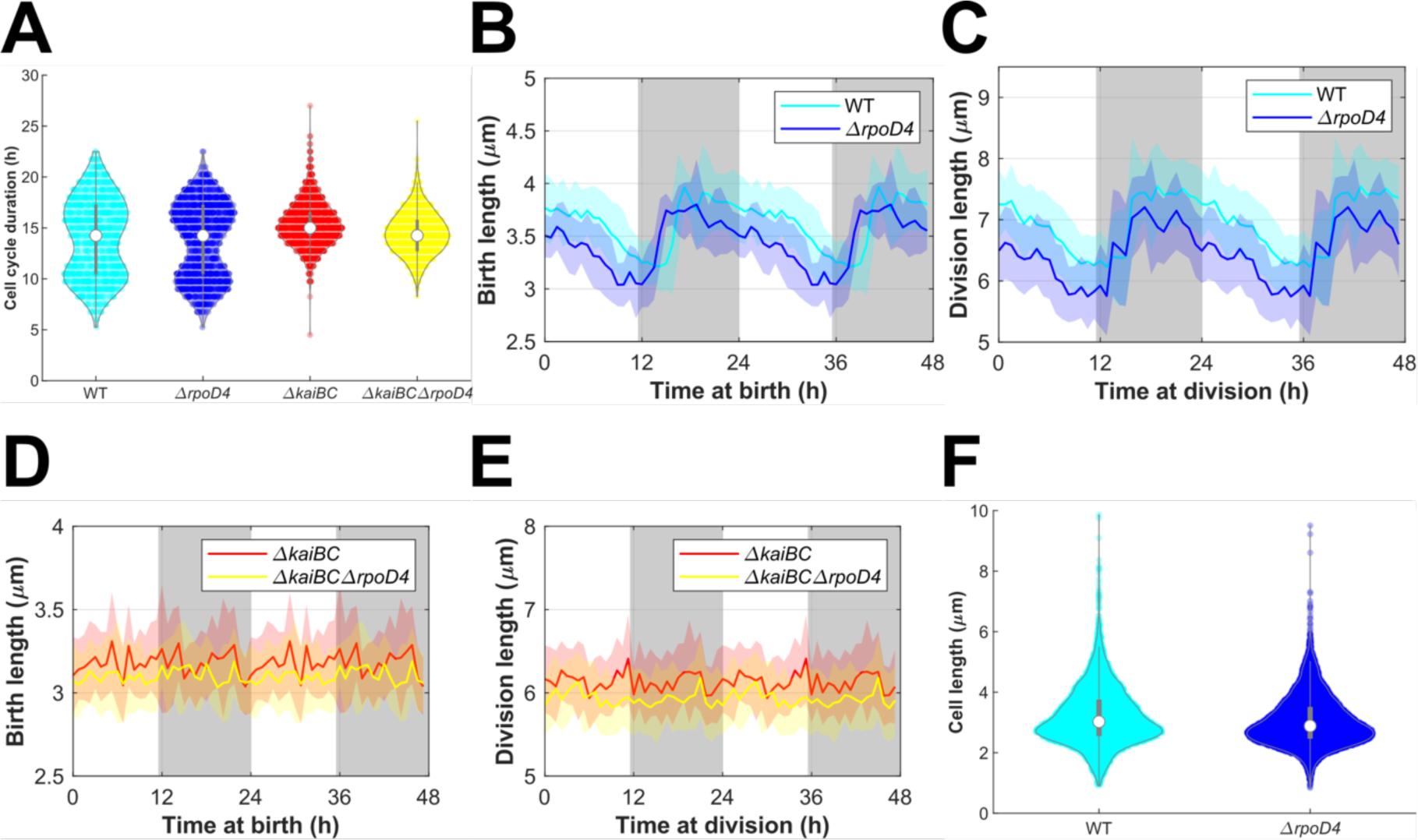
Deletion of *rpoD4* reduces length of cells grown on agarose pads and liquid cultures. (**A**) Violin plot of doubling time, (**B&D**) mean birth length plotted against time at birth, and (**C&E**) mean division length plotted against time at division of WT, *ΔrpoD4*, *ΔkaiBC* and *ΔrpoD4ΔkaiBC* strains carrying P*_rpoD4_*-EYFP-LVA reporters grown on agarose pads under constant light (c.a. 18 μE m^-2^ s^-1^). 1140 WT, 1310 *ΔrpoD4*, 1350 *ΔkaiBC*, and 1379 *ΔrpoD4ΔkaiBC* cells from two independent experiments were collected for analysis. All data are depicted for two cycles to highlight periodicity. The subjective day and night are indicated by the white and grey background shades. Cyan, blue, red and yellow shades represent one standard deviation from the mean. (**F**) Cell length distribution of mixed-phase WT (n = 6028) and *ΔrpoD4* (n = 8702) cells grown in liquid batch cultures. Samples were collected from two independent experiments when the cultures were at the mid-exponential phase (i.e., OD750 ∼ 0.5). The mean cell length of *ΔrpoD4* (3.042 ± 0.809 μm) is 4.72% smaller than WT (3.192 ± 0.957 μm). The cell size difference is statistically significant, but the size reduction effect of *ΔrpoD4* is small (p = 2.29e-22, Wilcoxon rank-sum test, df = 14728, ES = 0.172).

**Figure S6.**
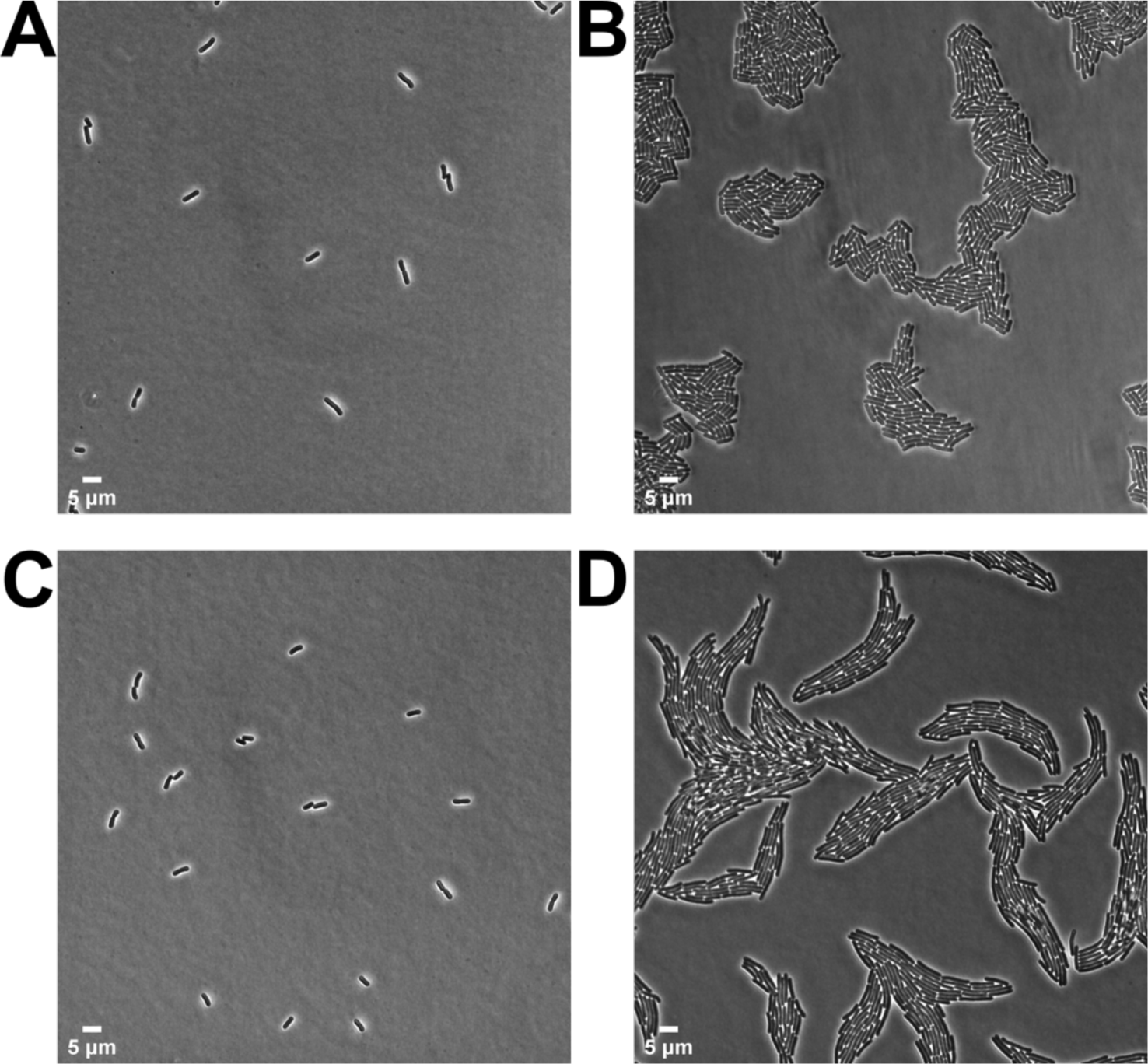
Overexpression of *rpoD4* increases size of cells grown on agarose pads. (**A**) First (t = 1.5 h after entrainment ends) and (**B**) last frames (t = 146.25 h) of a time-lapse movie of WT cells grown on an agarose pad. (**C**) First and (**D**) last frames of a time-lapse movie of P*_trc_*-*rpoD4* cells grown on an agarose pad supplemented with 100 μM IPTG.

**Figure S7.**
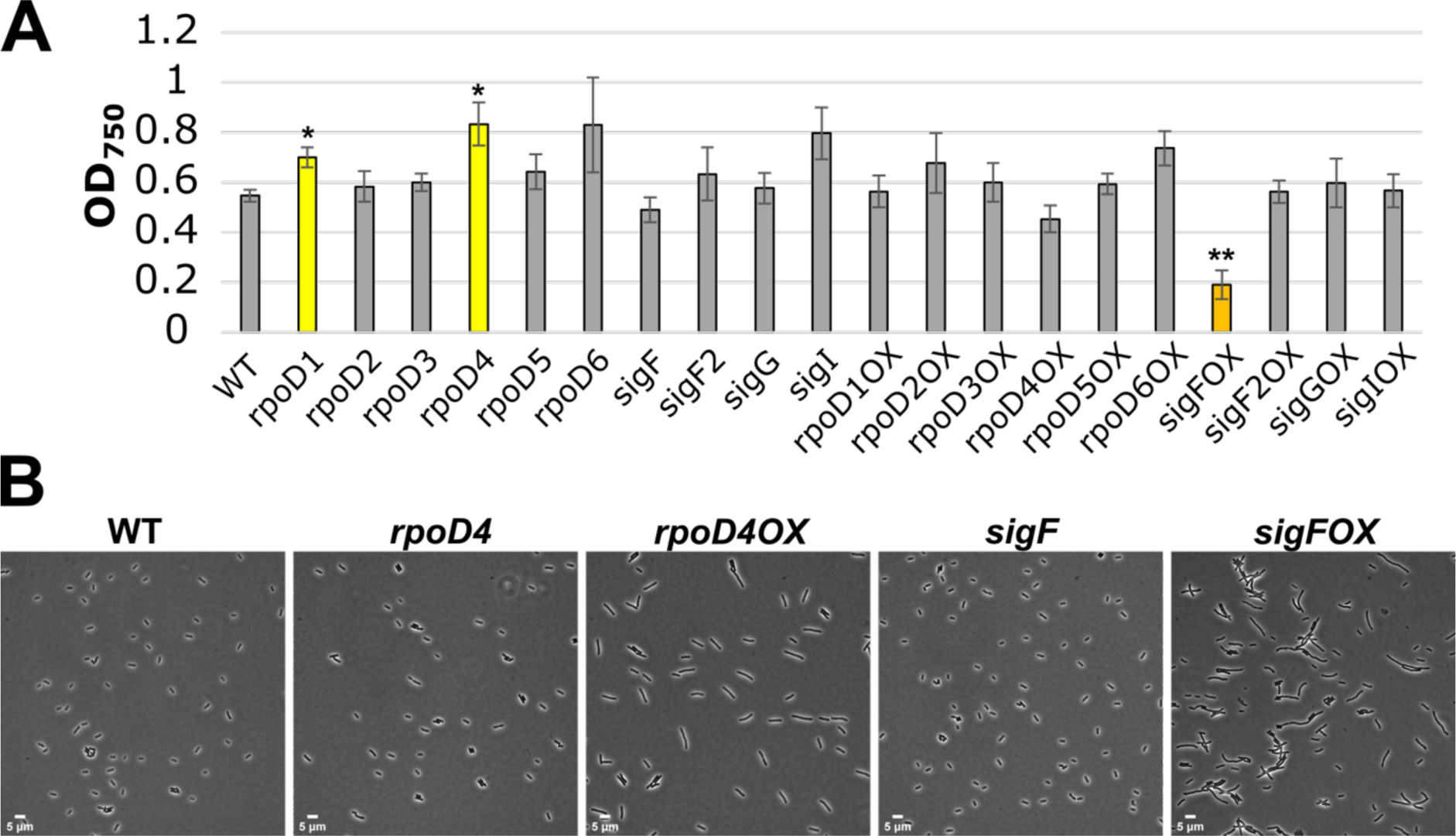
Overexpression of *rpoD4* does not affect cell growth in liquid cultures. (**A**) OD_750_ of WT and P*_trc_*-sigma incubated with or without 1 mM IPTG for 3 days. Data shown as mean ± SEM. Strains with significantly different OD_750_ compared to WT are highlighted with different colours (*: p<0.05 (yellow), **: p < 0.01 (orange)). Data were collected from three independent experiments (**B**) Snapshots of WT, P*_trc_*-*rpoD4*, P*_trc_*-*sigF* incubated with (*sigmaOX*) or without 1 mM IPTG for 3 days (all cultures were diluted to OD_750_ = 0.3 before snapshot).

**Figure S8.**
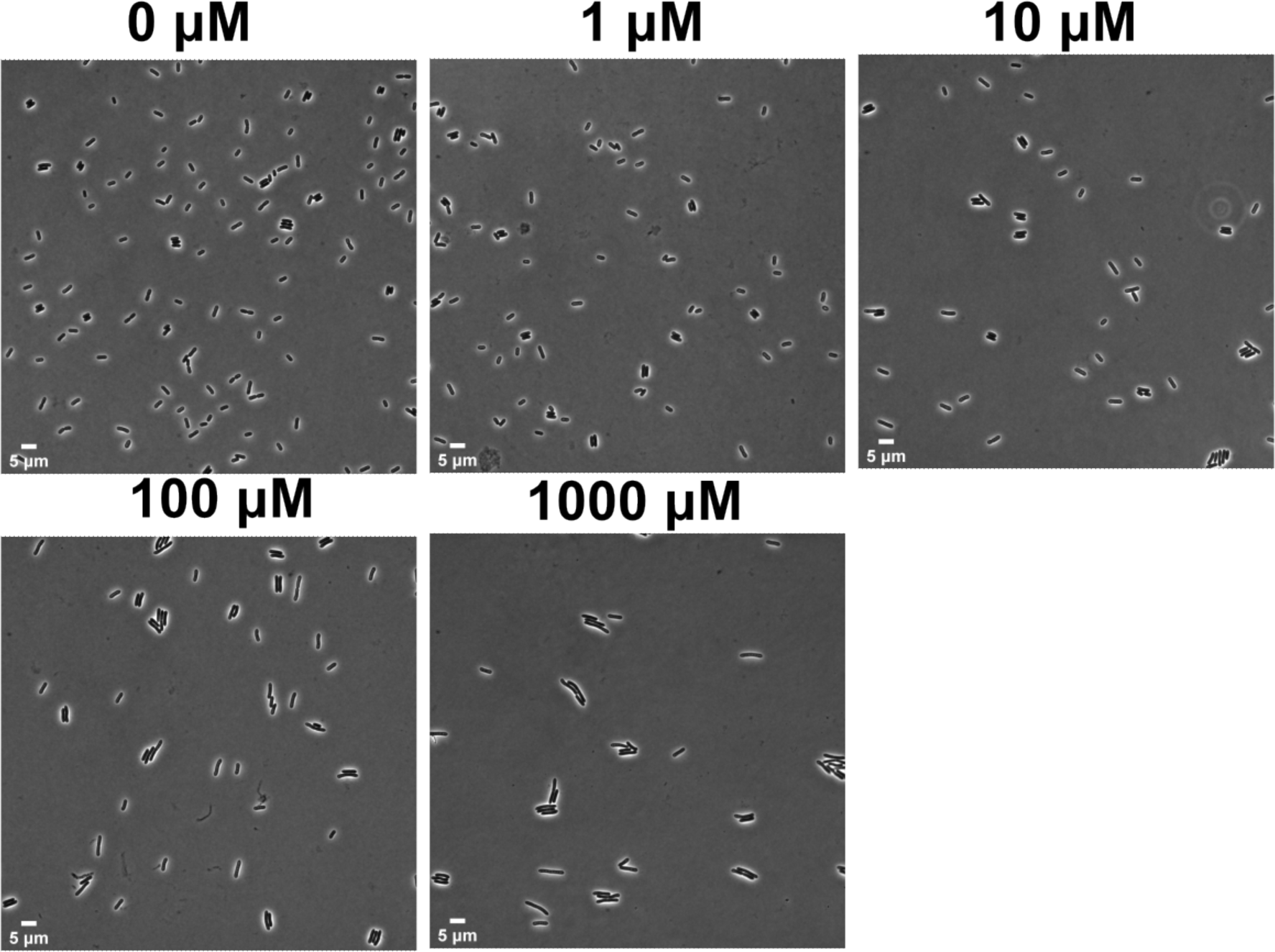
Dose-response relationship between the *rpoD4* expression level and cell size. Snapshots of P*_trc_*-*rpoD4* cells (starting OD_750_ = 0.05) incubated with 0, 1, 10, 100 and 1000 μM IPTG for 3 days under constant light (20-25 μE m^-2^ s^-1^) in batch cultures.

**Figure S9.**
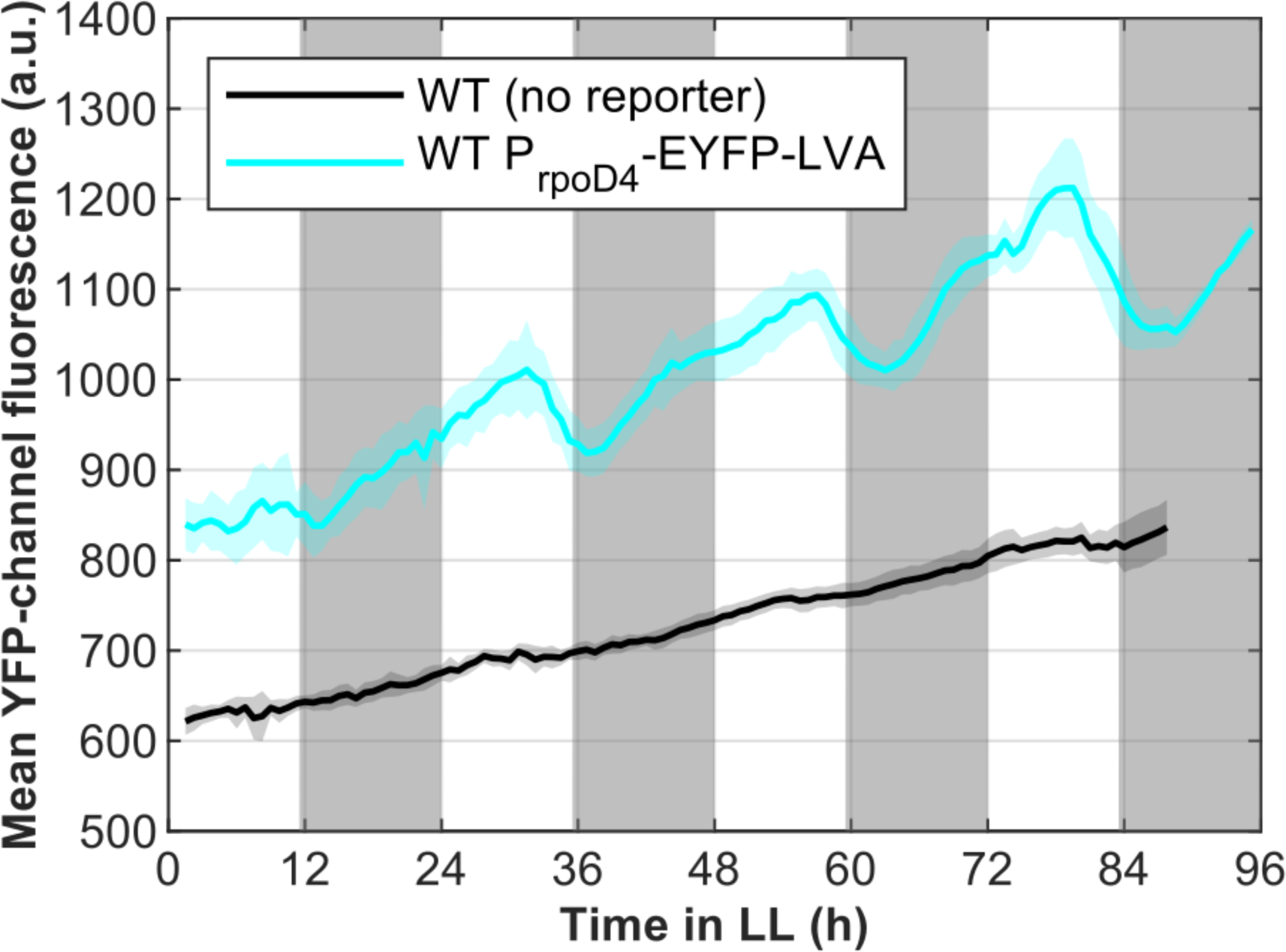
Autofluorescence of WT cells in YFP channel increases over time. Mean YFP channel fluorescence of WT cells with no reporter (black) and P*_rpoD4_*-EYFP-LVA cells (cyan) grown on agarose pads under constant light (c.a. 18 μE m^-2^ s^-1^). The solid lines represent the mean fluorescence averaged from 9 movies in 1 experiment. The shades represent one standard deviation from the mean.

## Bibliography

Andersson CR, Tsinoremas NF, Shelton J, Lebedeva N V., Yarrow J, Min H & Golden SS (2000) Application of bioluminescence to the study of circadian rhythms in cyanobacteria. Methods Enzymol. 305: 527–542

Belbin FE, Noordally ZB, Wetherill SJ, Atkins KA, Franklin KA & Dodd AN (2017) Integration of light and circadian signals that regulate chloroplast transcription by a nuclear-encoded sigma factor. New Phytol. 213: 727

Bertani G (1951) Studies on lysogenesis. I. The mode of phage liberation by lysogenic Escherichia coli. J. Bacteriol. 62: 293–300

Bervoets I, Van Brempt M, Van Nerom K, Van Hove B, Maertens J, De Mey M & Charlier D (2018) A sigma factor toolbox for orthogonal gene expression in Escherichia coli. Nucleic Acids Res. 46: 2133

Bieler J, Cannavo R, Gustafson K, Gobet C, Gatfield D & Naef F (2014) Robust synchronization of coupled circadian and cell cycle oscillators in single mammalian cells. Mol. Syst. Biol. 10: 739

Chabot JR, Pedraza JM, Luitel P & van Oudenaarden A (2007) Stochastic gene expression out-of-steady-state in the cyanobacterial circadian clock. Nature 450: 1249–1252

Cohen SE & Golden SS (2015) Circadian Rhythms in Cyanobacteria. Microbiol. Mol. Biol. Rev. 79: 373–385

Cuitun-Coronado D & Dodd AN (2020) Plant Sigma Factors. eLS: 1–9

Ditty JL, Canales SR, Anderson BE, Williams SB & Golden SS (2005) Stability of the Synechococcus elongatus PCC 7942 circadian clock under directed anti-phase expression of the kai genes. Microbiology 151: 2605–2613

Doherty CJ & Kay SA (2010) Circadian Control of Global Gene Expression Patterns. Annu Rev Genet 44: 419–444

Dong G, Yang Q, Wang Q, Kim YI, Wood TL, Osteryoung KW, van Oudenaarden A & Golden SS (2010) Elevated ATPase Activity of KaiC Applies a Circadian Checkpoint on Cell Division in Synechococcus elongatus. Cell 140: 529–539

Feillet C, van der Horst GTJ, Levi F, Rand DA & Delaunay F (2015) Coupling between the circadian clock and cell cycle oscillators: Implication for healthy cells and malignant growth. Front. Neurol. 6: 1–7

Fleming KE & O’Shea EK (2018) An RpaA-Dependent Sigma Factor Cascade Sets the Timing of Circadian Transcriptional Rhythms in Synechococcus elongatus. Cell Rep. 25: 2937–2945.e3

Gan S & O’Shea EK (2017) An Unstable Singularity Underlies Stochastic Phasing of the Circadian Clock in Individual Cyanobacterial Cells. Mol. Cell 67: 659–672.e12

Gibson DG, Young L, Chuang R, Venter JC, Hutchison C a & Smith HO (2009) Enzymatic assembly of DNA molecules up to several hundred kilobases. Nat. Methods 6: 343–5

Golden SS & Canales SR (2003) Cyanobacterial circadian clocks — timing is everything. Nat. Rev. Microbiol. 1: 191–199

Gould PD, Domijan M, Greenwood M, Tokuda IT, Rees H, Kozma-Bognar L, Hall AJW & Locke JCW (2018) Coordination of robust single cell rhythms in the Arabidopsis circadian clock via spatial waves of gene expression. eLife 7: 1–20

Hanaoka M, Takai N, Hosokawa N, Fujiwara M, Akimoto Y, Kobori N, Iwasaki H, Kondo T & Tanaka K (2012) RpaB, another response regulator operating circadian clock-dependent transcriptional regulation in Synechococcus elongatus PCC 7942. J. Biol. Chem. 287: 26321–26327

Hanaoka M & Tanaka K (2008) Dynamics of RpaB-promoter interaction during high light stress, revealed by chromatin immunoprecipitation (ChIP) analysis in Synechococcus elongatus PCC 7942. Plant J. 56: 327–335

Hurley JM, Loros JJ & Dunlap JC (2016) Circadian Oscillators: Around the Transcription-Translation Feedback Loop and on to Output. Trends Biochem. Sci. 41: 834

Hutchison AL, Maienschein-Cline M, Chiang AH, Tabei SMA, Gudjonson H, Bahroos N, Allada R & Dinner AR (2015) Improved Statistical Methods Enable Greater Sensitivity in Rhythm Detection for Genome-Wide Data. PLOS Comput. Biol. 11: e1004094

Irwin CR, Farmer A, Willer DO & Evans DH (2012) In-fusion® cloning with vaccinia virus DNA polymerase. Methods Mol. Biol. 890: 23–35

Ito H, Mutsuda M, Murayama Y, Tomita J, Hosokawa N, Terauchi K, Sugita C, Sugita M, Kondo T & Iwasaki H (2009) Cyanobacterial daily life with Kai-based circadian and diurnal genome-wide transcriptional control in Synechococcus elongatus. Proc. Natl. Acad. Sci. U. S. A. 106: 14168–14173

Ivleva NB, Bramlett MR, Lindahl PA & Golden SS (2005) LdpA: A component of the circadian clock senses redox state of the cell. EMBO J.

Johnson CH & Egli M (2014) Metabolic compensation and circadian resilience in prokaryotic cyanobacteria. Annu. Rev. Biochem. 83: 221–247

Jun S & Rust MJ (2017) A Fundamental Unit of Cell Size in Bacteria. Trends Genet. 33: 433–435

Kobayashi I, Watanabe S, Kanesaki Y, Shimada T, Yoshikawa H & Tanaka K (2017) Conserved two-component Hik34-Rre1 module directly activates heat-stress inducible transcription of major chaperone and other genes in Synechococcus elongatus PCC 7942. Mol. Microbiol. 104: 260–277

Kulkarni RD & Golden SS (1997) mRNA stability is regulated by a coding-region element and the unique 5’ untranslated leader sequences of the three Synechococcus psbA transcripts. Mol. Microbiol. 24: 1131–1142

Lambert G, Chew J & Rust MJ (2016) Costs of Clock-Environment Misalignment in Individual Cyanobacterial Cells. Biophys. J.

Liao Y & Rust MJ (2021) The circadian clock ensures successful DNA replication in cyanobacteria. Proc. Natl. Acad. Sci. U. S. A. 118: 1–8

Mackey SR, Ditty JL, Clerico EM & Golden SS (2007) Detection of rhythmic bioluminescence from luciferase reporters in cyanobacteria. Methods Mol. Biol. 362: 115–129

Markson JS, Piechura JR, Puszynska AM & O’Shea EK (2013) Circadian control of global gene expression by the cyanobacterial master regulator RpaA. Cell 155: 1396–1408

Martins BM, Das AK, Antunes L & Locke JC (2016) Frequency doubling in the cyanobacterial circadian clock. Mol. Syst. Biol. 12: 896

Martins BMC, Tooke AK, Thomas P & Locke JCW (2018) Cell size control driven by the circadian clock and environment in cyanobacteria. Proc. Natl. Acad. Sci. U. S. A. 155(48): E11415–E11424

Mori T, Binder B & Johnson CH (1996) Circadian gating of cell division in cyanobacteria growing with average doubling times of less than 24 hours. Proc. Natl. Acad. Sci. U. S. A. 93: 10183–10188

Mori T & Johnson CH (2001) Independence of circadian timing from cell division in cyanobacteria. J. Bacteriol. 183: 2439–2444

Moronta-Barrios F, Espinosa J & Contreras A (2012) In vivo features of signal transduction by the essential response regulator RpaB from Synechococcus elongatus PCC 7942. Microbiology 158: 1229–1237

Nair U, Ditty JL, Min H & Golden SS (2002) Roles for Sigma Factors in Global Circadian Regulation of the Cyanobacterial Genome Roles for Sigma Factors in Global Circadian Regulation of the Cyanobacterial Genome. 184: 3530–3538

Noordally ZB, Ishii K, Atkins KA, Wetherill SJ, Kusakina J, Walton EJ, Kato M, Azuma M, Tanaka K, Hanaoka M & Dodd AN (2013) Circadian control of chloroplast transcription by a nuclear-encoded timing signal. Science 339: 1316–1319

Paijmans J, Bosman M, Ten Wolde PR & Lubensky DK (2016) Discrete gene replication events drive coupling between the cell cycle and circadian clocks. Proc. Natl. Acad. Sci. U. S. A. 113: 4063–4068

Paranjpe DA & Sharma VK (2005) Evolution of temporal order in living organisms. J. Circadian Rhythms 3(1): 7

Pattanayak G & Rust MJ (2014) The cyanobacterial clock and metabolism. Curr. Opin. Microbiol. 18: 90–95

Pattanayak GK, Lambert G, Bernat K & Rust MJ (2015) Controlling the Cyanobacterial Clock by Synthetically Rewiring Metabolism. Cell Rep. 13: 2362–2367

Plautz JD, Straume M, Stanewsky R, Jamison CF, Brandes C, Dowse HB, Hall JC & Kay SA (1997) Quantitative analysis of Drosophila period gene transcription in living animals. J. Biol. Rhythms 12: 204–217

Schwall CP, Loman TE, Martins BMC, Cortijo S, Villava C, Kusmartsev V, Livesey T, Saez T & Locke JCW (2021) Tunable phenotypic variability through an autoregulatory alternative sigma factor circuit. Mol. Syst. Biol. 17: 1–16

Seki A, Hanaoka M, Akimoto Y, Masuda S, Iwasaki H & Tanaka K (2007) Induction of a group 2 σ factor, RPOD3, by high light and the underlying mechanism in Synechococcus elongatus PCC 7942. J. Biol. Chem. 282: 36887–36894

Teng S-W, Mukherji S, Moffitt JR, de Buyl S & O’Shea EK (2013) Robust circadian oscillations in growing cyanobacteria require transcriptional feedback. Science 340: 737–40

Vijayan V, Zuzow R & O’Shea EK (2009) Oscillations in supercoiling drive circadian gene expression in cyanobacteria. Proc. Natl. Acad. Sci. U. S. A. 106: 22564–8

Welkie DG, Rubin BE, Chang YG, Diamond S, Rifkin SA, LiWang A & Golden SS (2018) Genome-wide fitness assessment during diurnal growth reveals an expanded role of the cyanobacterial circadian clock protein KaiA. Proc. Natl. Acad. Sci. U. S. A. 115: E7174–E7183

Yang Q, Pando BF, Dong G, Golden SS & Van Oudenaarden A (2010) Circadian gating of the cell cycle revealed in single cyanobacterial cells. Science. 327: 1522–1526

Yang Y, Lam V, Adomako M, Simkovsky R, Jakob A, Rockwell NC, Cohen SE, Taton A, Wang J, Clark Lagarias J, Wilde A, Nobles DR, Brand JJ & Golden SS (2018) Phototaxis in a wild isolate of the cyanobacterium Synechococcus elongatus. Proc. Natl. Acad. Sci. U. S. A.

Young JW, Locke JCW, Altinok A, Rosenfeld N, Bacarian T, Swain PS, Mjolsness E & Elowitz MB (2012) Measuring single-cell gene expression dynamics in bacteria using fluorescence time-lapse microscopy. Nat. Protoc. 7: 80–8

Zheng X yu & O’Shea EK (2017) Cyanobacteria Maintain Constant Protein Concentration despite Genome Copy-Number Variation. Cell Rep. 19: 497–504

Zielinski T, Moore AM, Troup E, Halliday KJ & Millar AJ (2014) Strengths and limitations of period estimation methods for circadian data. PLoS One. 9(5): e96462

